# Single-cell transcriptomics unveils xylem cell development and evolution

**DOI:** 10.1101/2021.10.16.464627

**Authors:** Chia-Chun Tung, Shang-Che Kuo, Chia-Ling Yang, Chia-En Huang, Jhong-He Yu, Ying-Hsuan Sun, Peng Shuai, Jung-Chen Su, Chuan Ku, Ying-Chung Jimmy Lin

**Author notes:** These authors contributed equally (Chia-Chun Tung, Shang-Che Kuo, Chia-Ling Yang).

## Abstract

As the most abundant tissue on Earth^1^, xylem is responsible for lateral growth in plants. Typical xylem has a radial system composed of ray parenchyma cells and an axial system of fusiform cells^2^. In most angiosperms, fusiform cells are a combination of vessel elements for water transportation and libriform fibers for mechanical support, while both functions are performed together by tracheids in other vascular plants^2^. However, little is known about the developmental programs and evolutionary relationships of these xylem cell types. Through both single-cell and laser-capture microdissection transcriptomic profiling, here we demonstrate the developmental lineages of ray and fusiform cells in stem-differentiating xylem across four divergent woody angiosperms. Cross-species analyses of single-cell trajectories reveal highly conserved ray, yet variable fusiform, lineages across angiosperms. Core eudicots *Populus trichocarpa* and *Eucalyptus grandis* share nearly identical fusiform lineages. The tracheids in the basal eudicot *Trochodendron aralioides*, an evolutionarily reversed character^3, 4^, exhibit strong transcriptomic similarity to vessel elements but not libriform fibers, suggesting that water transportation, instead of mechanical support, is the major feature. We also found that the more basal angiosperm *Liriodendron chinense* has a fusiform lineage distinct from that in core eudicots. This evo-developmental framework provides a comprehensive understanding of the formation of xylem cell lineages across multiple plant species spanning over a hundred million years of evolutionary history^5^.

## Main

Plant development involves apical and lateral growth. Apical growth elongates the main plant axes by growing upward from shoot apical meristem (SAM) into shoots, leaves or flowers and downward through root apical meristem (RAM) into roots^6^. Lateral growth, which thickens the plant body, is bifacial in nature with a single layer of initials in vascular cambium growing inward into xylem and outward into phloem^7–10^. The developmental trajectories of apical roots and shoots have been explored using single-cell RNA sequencing (scRNA-seq)^11–22^. The spatial and temporal distribution of individual cells from different stages reveals highly heterogeneous transcriptomes of cell lineages during proliferation, identity determination, and differentiation. Isolation of single protoplasts has allowed high-resolution transcriptomic analyses of leaves, flowers and roots. These organs tend to yield few cells from vascular tissue due to their relatively small proportion of vascular cells^11–22^, limiting our understanding of the molecular mechanisms in xylem development at the tissue level. The developmental trajectories of different cell types in xylem, especially secondary xylem, still remain unknown.

Xylem comprises axial and radial systems that develop from two types of initials in vascular cambium^2^. The axial system differentiated from fusiform initials is represented as a combination of vessel elements and libriform fibers in most angiosperms and as tracheids in other vascular plants. The radial system contains ray parenchyma cells from ray initials. Over the past hundred years, the development of xylem cell types has mostly been studied through anatomical observations, with the molecular mechanisms behind their differentiation still largely unknown. Here we used scRNA-seq on the stem-differentiating xylem (SDX) of four representative woody species with distinct cell types, including tracheids, vessel elements, libriform fibers and ray parenchyma cells, from core eudicots, basal eudicots and magnoliids of angiosperms to comprehensively explore xylem development in an evolutionary context.

### Single-cell and cell-type RNA sequencing reveal SDX cell identities

We profile the transcriptomes of 4,705 individual SDX cells from *P. trichocarpa* (black cottonwood) using 10x Genomics Chromium technology (Fig. 1a). Unsupervised K-means clustering divides the cells into 10 clusters Ptr1 to Ptr10 (Extended Data Fig. 1a), which are visualized using Uniform Manifold Approximation and Projection (UMAP) (Fig. 1b). Current definitions of SDX cell types are mainly based on anatomical observations^2, 23^, and no suitable marker genes are available for assigning cell types to the scRNA-seq cell clusters. To correlate morphology and transcriptomes, we established strategies of laser capture microdissection (LCM) to sequence the mRNA (lcmRNA-seq) of three cell types separately by preventing cross contamination. Transverse sections are used to collect libriform fibers (Extended Data Fig. 2), and the laser cutting pathway avoids the locations of vessel elements and ray parenchyma cells (red area in Fig. 1c-e, Extended Data Fig. 3). Tangential sections are used for collecting vessel elements and ray parenchyma cells (Fig. 1g–i, k–m, Extended Data Figs. 2, 4). For vessel elements, we only cut the cells with obvious and intact pitted cell wall (blue area in Fig. 1g–i, Extended Data Fig. 4a–c). For ray parenchyma cells, we only collect the cells located at the center of each tangential section (pink area in Fig. 1k–m, Extended Data Fig. 4d–h), avoiding the ray parenchyma cells on both sides that could not be easily separated from neighbor cells (Fig. 1k–m).

**Fig. 1.**
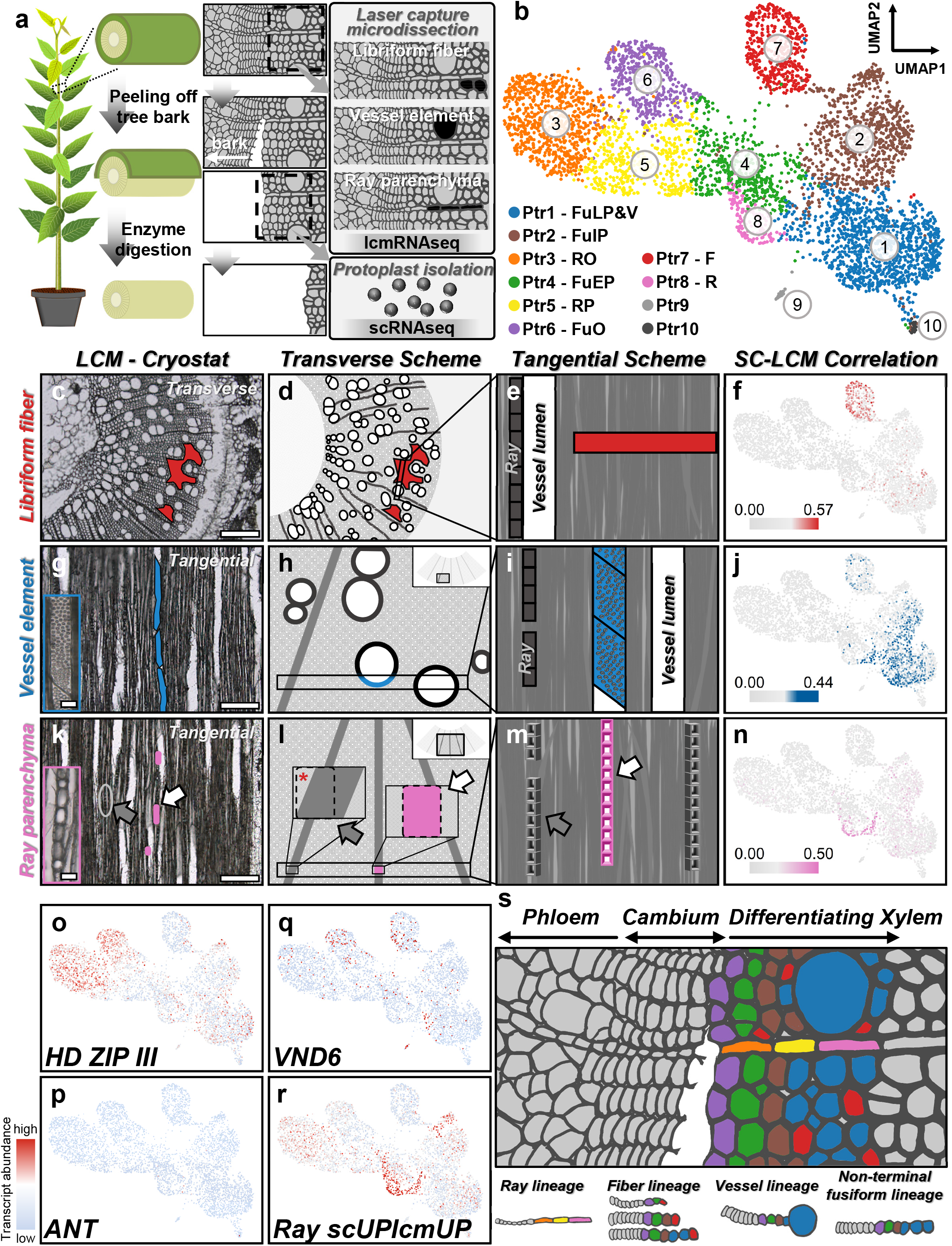
scRNA-seq and lcmRNA-seq reveal cell clusters involved in SDX development. **a**, Schematic of the workflow for lcmRNA-seq of libriform fibers, vessel elements and ray parenchyma cells (shown as black area) and scRNA-seq of SDX protoplasts. For SDX protoplast isolation, the stems of greenhouse-grown *P. trichocarpa* were processed by bark peeling and cell-wall enzyme digestion. **b**, Ten cell clusters, Ptr1 to Ptr10, were obtained through unsupervised K-means clustering and visualized by UMAP. FuLP&V, fusiform late precursor and vessel element. FuIP, fusiform intermediate precursor. RO, ray organizer. FuEP, fusiform early precursor. RP, ray precursor. FuO, fusiform organizer. F, libriform fiber. R, ray parenchyma cell. **c**–**n**, LCM for three cell type collection and their transcriptomic correlation to scRNA-seq results. **c**, **g**, **k**, Real transverse or tangential sections of *P. trichocarpa* stems for libriform fiber (red area) (**c**), vessel element (blue area) (**g**) and ray parenchyma cell (pink area) (**k**) collection. Scale bars, 200 µm. The blue and pink rectangles at the left bottom in **g** and **k** are the magnified vessel elements and ray parenchyma cells. Scale bars, 25 µm. **d**, **e**, **h**, **i**, **l**, **m**, The illustrations for three cell type collection from transverse and tangential perspectives. **e**, **i**, The white box represents a vessel element with empty lumen. **l**, The pink box with dashed lines is the area for ray parenchyma cell collection. The other dashed-line box indicates the potential contamination caused by collecting the ray parenchyma cells not from the middle of a section. Such dashed-line box contains dark-grey area as ray parenchyma cells and light-grey area as the neighbor libriform fibers shown as the red asterisk. **k**, **l**, **m**, The white and grey arrows indicate the collected and avoided ray parenchyma cells, respectively. **f**, **j**, **n**, The transcriptomic correlations of libriform fibers (**f**), vessel elements (**j**) and ray parenchyma cells (**n**) to the single-cell transcriptomes, respectively. **o**–**r**, The transcript abundance of *HD ZIP III* (**o**), *ANT* (**p**), *VND6* (**q**) and scUPlcmUP genes in ray parenchyma cells (**r**) in scRNA-seq results. **s**, The proposed cell lineages during SDX development shown on a schematic transverse section of *P. trichocarpa*, including ray, fiber, vessel lineages and a non-terminal fusiform lineage. The SDX protoplast isolation releases around six layers of SDX cells from the stem surface after debarking, so around six layers are labeled with colors. The phloem part removed with tree bark and the inner part (inner than six layers) are labeled as grey.

To identify the cell types of the scRNA-seq cell clusters, we examine the transcriptomic correlation between each individual cell from scRNA-seq and each cell type from lcmRNA-seq. The cell clusters Ptr7 (red), Ptr1 (blue) and Ptr8 (pink) show the highest correlation to vessel elements, libriform fibers and ray parenchyma cells, respectively (Fig. 1f, j, n, Extended Data Fig. 5). There are also the highest numbers of differentially expressed genes (DEGs) up-regulated in both these cell clusters and their corresponding lcmRNA-seq cell type transcriptomes (scUPlcmUP) (Extended Data Fig. 6).

We found that the scUPlcmUP genes of libriform fibers (Ptr7) are heavily enriched with secondary cell wall (SCW) biosynthesis genes, especially those for monolignol biosynthesis (Extended Data Figs. 7a–n, 8). Ray parenchyma cells also have moderate expression of monolignol biosynthesis genes (Extended Data Fig. 8), consistent with previous observations that parenchyma cells in *Arabidopsis* and *Picea* also contribute to lignification^24–26^. Vessel elements barely express monolignol biosynthesis genes (Extended Data Fig. 8), supporting the good-neighbor or post-mortem hypothesis^27, 28^ that the lignification of vessel elements is non-cell-autonomous. Expansins are proteins that loosen cell wall to promote cell expansion^29, 30^. The most highly expressed expansin gene in SDX of *P. trichocarpa*, *Potri.001G240900* (Extended Data Fig. 9), is significantly enriched in vessel elements (Extended Data Fig. 7o). Since vessel elements have the widest cell lumen among three cell types, the highest expansin transcript abundance would facilitate the enormous requirement for radial cell wall expansion of vessel elements. Furthermore, we found many photosynthesis-related genes among the scUPlcmUP genes in ray parenchyma cells (Extended Data Fig.7p–w), which agrees with previous observations that ray parenchyma cells possess chlorophyll-containing plastids to carry out woody tissue photosynthesis^31–34^. By integrating scRNA-seq–lcmRNA-seq correlation, scUPlcmUP gene distribution and known gene functions, we revealed the transcriptomic profiles and single-cell clusters of vessel elements, libriform fibers and ray parenchyma cells.

### Two distinct lineages of fusiform and ray cell differentiation from organizer cells

The three cell types at terminal differentiated stages are all localized in the right part of the UMAP plot (Fig. 1b), leaving the rest of the cell clusters unidentified (Fig. 1b). All stem cells from SAM, RAM and vascular cambium are constantly maintained by the organizer cells^6, 35^. The HD ZIP III family transcription factor in *Arabidopsis* (AtHB8) is sufficient to induce the function of organizer cells derived from vascular cambium to determine xylem identity^10^, and was also reported to participate in interfascicular cambium development in *Populus*^36^. The homologous genes of *AtHB8* in *P. trichocarpa* were mainly expressed in Ptr3 and Ptr6 (Fig. 1o, Extended Data Fig. 10a). We did not detect the expression of *ANT* (a stem cell/initial cell marker gene^37^; Fig. 1p, Extended Data Fig. 10b) or *PEAR1* (a phloem marker gene^37^; Extended Data Fig. 10c) in any of the cell clusters, as expected from the removal of initials and phloem during the debarking step of protoplast preparation (Fig. 1a). It suggests the position of organizer cells in SDX (Fig. 1s) is similar to that of SAM organizer cells between stem cells/initials and xylem cells^6, 35^.

From previous anatomical studies, SDX development is a combination of two trajectories of fusiform and ray initials^2, 23^. The two cell clusters Ptr3 and Ptr6 likely represent two types of organizer cells (Fig. 1b). Ray initials are generated from the asymmetric cell division of fusiform initials^2, 23, 38, 39^. We observed some fusiform and ray organizer cells intermixed with each other (Fig. 1b), demonstrating that the ray organizer cells divided from ray initials still retained high transcriptomic similarity to fusiform organizer cells. The transcript abundance of the respective up-regulated DEGs of vessel elements (Ptr1) and ray parenchyma cells (Ptr8) (Extended Data Fig. 11) further resolved the cell clusters into two continuous cell lineages. The sequence Ptr6-Ptr4-Ptr2-Ptr1 is the developmental path from fusiform organizer (Ptr6) through fusiform early precursor (Ptr4) and fusiform intermediate precursor (Ptr2) to vessel elements (Ptr1). *VND6* members in the *VNS* family, the master regulators of vessel element differentiation^40^, and the aforementioned expansin gene (*Potri.001G240900*) are also expressed in this fusiform cell lineage (Fig. 1q, Extended Data Figs. 10g, 11a). The cell clusters Ptr3 (ray organizer), Ptr5 (ray precursor) and Ptr8 (ray parenchyma) form the other cell lineage (Fig. 1b, Extended Data Fig. 11b), which is derived from ray initials.

We notice that fusiform and ray lineages converge at fusiform early precursors (Ptr4) and ray precursors (Ptr5) before entering the terminal differentiated stages (Fig. 1b). To validate this observation, we applied another cell clustering and dimensionality reduction pipeline, MetaCell^41^, to the scRNA-seq dataset. The MetaCell plot similarly shows that these two cell lineages merge together at the intermediate stage of differentiation (Extended Data Fig. 12). Among all pairwise comparisons of cell cluster transcriptomic profiles, the pair Ptr4-Ptr5 also has the highest correlation (Extended Data Fig. 13). This striking transcriptomic similarity between fusiform early precursors and ray precursors suggests that fusiform and ray cells would differentiate into a similar state before continuing their unique developmental pathways.

It is still debated whether the cell fate of fusiform cells is determined at the very beginning in the initial cells or later during the differentiation of xylary derivatives^8^. Based on the ubiquitous expression of *VNS* family genes in all early fusiform cells^42^, it has been proposed that all fusiform cells originally differentiate toward vessel elements but become committed toward libriform fibers upon an external or positional cue^8^. As a response to mechanical stress, for example, the formation of tension wood contains drastically increased libriform fibers^2, 43^. A closer look at the fusiform cell clusters reveals that the early (Ptr4) and intermediate (Ptr2) fusiform precursors and Ptr1 all have cells intermixed with libriform fibers (Ptr7) (Extended Data Fig. 14a), suggesting that these cell types, including Ptr1, might have the potency to transit into libriform fibers (Extended Data Fig. 14b). The results indicate that Ptr1 is a mixture of terminal differentiated vessel elements and fusiform cells differentiating toward vessel elements. Ptr1 thus represents vessel element/late fusiform precursor. Interestingly, the libriform fibers (Ptr7) are the most disjoint cell cluster (Fig. 1b, Extended Data Fig. 12), implying the differentiation into libriform fibers took place rapidly, leaving few intermediate cells to be captured for scRNA-seq. Such disconnection in cell states has also been demonstrated for malaria parasite cell types, where an abrupt transcriptomic change is caused by fast turn-on/off of transcriptional modules^44^. Our results suggest that fusiform cells differentiate into vessel elements by default, which can be rapidly switched to libriform fibers by external signals.

In summary, we uncovered the detailed cell lineages in *P. trichocarpa* SDX (Extended Data Fig. 15). Through the clustering of single-cell transcriptomic profiles and identification of different cell types, we provide novel marker genes (Supplementary Table 1) with expression specific to each cell type (Extended Data Figs. 16, 17). Using a conserved developmental mechanism involving stem cells and organizer cells comparable to those in SAM and RAM, xylem fusiform and ray cells exert their differentiation through two distinct trajectories.

### Identifying SDX cell types across angiosperms

To further explore SDX development in other woody plants, we performed scRNA-seq of SDX cells in another core eudicot *E. grandis*, a basal eudicot *T. aralioides* (wheel tree) and a magnoliid *L. chinense* (Chinese tulip poplar). In contrast to the vast majority of flowering plants, *T. aralioides* possesses tracheids instead of vessel elements and libriform fibers, which is a rare trait reversal during angiosperm evolution (Fig. 2a)^3, 4^. *L*. *chinense* protoplasts were sequenced using the 10x Genomics platform as for *P*. *trichocarpa*. Samples from *E*. *grandis* and field-collected *T*. *aralioides* were replete with cellular debris, which prompted us to adopt fluorescence-activated cell sorting coupled with the plate-based MARS-seq2.0 protocol to sequence transcriptomes of single sorted protoplasts with minimal debris contamination. We obtained the transcriptomic profiles from 5494, 1993 and 2977 individual SDX cells, respectively, which were grouped into 10 cell clusters in each species. (Fig. 2b–d, Extended Data Fig. 1). The DEGs of each cell cluster in the three species were then identified (Supplementary Tables 2–4).

**Fig. 2.**
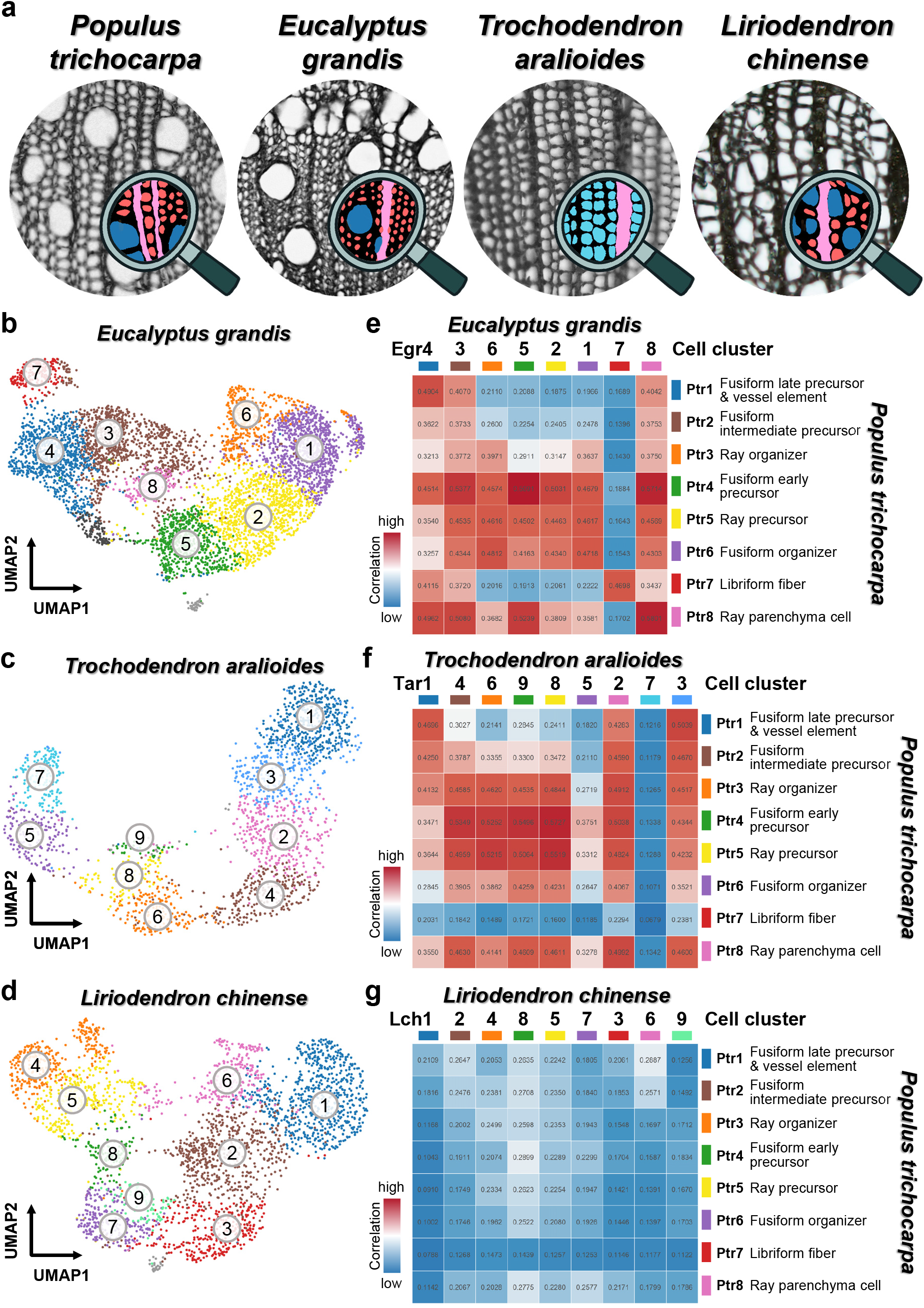
Comparative anatomy and single-cell transcriptomics of SDX in *P. trichocarpa*, *E. grandis*, *T. aralioides* and *L. chinense*. **a**, The SDX anatomy of four woody species. *P. trichocarpa*, *E. grandis* and *L. chinense* contain libriform fibers (red), vessel elements (blue) and ray parenchyma cells (pink). *T. aralioides* possesses tracheids (cyan) and ray parenchyma cells (pink). **b**–**d**, The unsupervised K-means clustering and UMAP plots from the SDX scRNA-seq results of *E. grandis* (**b**), *T. aralioides* (**c**) and *L. chinense* (**d**), respectively. **e**–**g**, The pairwise correlation analysis of the SDX scRNA-seq results between *P. trichocarpa* and *E. grandis* (**e**), *T. aralioides* (**f**) or *L. chinense* (**g**), respectively. The order of cell clusters in *E. grandis* (e.g. Egr4-3-6-5-2-1-7-8), *T. aralioides* or *L. chinense* are corresponding to those in Extended Data Fig. 1.

Cross-species pairwise correlation analysis of transcriptomic profiles was performed to compare the major cell clusters of *E. grandis*, *T. aralioides* and *L. chinense* to those of *P. trichocarpa* (Fig. 2e–g). Overall the correlations between *P. trichocarpa* and *E. grandis* or *T. aralioides* are higher than those between *P. trichocarpa* and *L. chinense* (Fig. 2e–g), reflecting the closer phylogenetic relationships among the three eudicots than between eudicots and the more basal magnoliid angiosperm. Considering that the SDX cell types in *L. chinense* are the same as in *P*. *trichocarpa*^45^ (Fig. 2a), two scenarios might explain their low transcriptomic correlation. First, the SDX cell differentiation might be regulated by only a few master regulators, so that despite the overall dissimilar transcriptomic profiles, the organizer cells still develop into the same cell types under the control of these master regulators. Second, the cell types are actually of different developmental origins but share similar morphology due to convergent evolution.

In general, we are able to identify many representative cell clusters in *E. grandis* corresponding to different cell types in *P. trichocarpa*. Egr5, Egr7 and Egr8 resembled the fusiform early precursors (Ptr4), libriform fibers (Ptr7) and ray parenchyma (Ptr8) cell clusters in *P. trichocarpa*, respectively, with the mutually highest correlation (mutual best-hit) in all-against-all comparison (Fig. 2e). Egr4 is the best-hit of vessel elements/fusiform late precursors (Ptr1), and Ptr1 is the second best-hit of Egr4 (Fig. 2e). The fusiform and ray organizer cells (Ptr6 and Ptr3) have a number of intermixed cells (Fig. 1b), possibly reflecting the derivation of ray initials from fusiform initials^2, 23, 38, 39^. Both Egr6 and Egr1 resemble fusiform organizer cells (Ptr6), indicating they could be the fusiform and ray organizer cells.

In *T. aralioides*, the cell clusters Tar1 and Tar3 show the highest correlations with *P*. *trichocarpa* fusiform late precursors/vessel elements (Ptr1) (Fig. 2f). As an evolutionarily reversed character, the terminally differentiated fusiform cells in *T. aralioides* are tracheids, which is a primitive form with the combined functions of vessel elements and libriform fibers. Tar1 and Tar3 clusters thus likely represent tracheids. Tar2 and ray parenchyma cells (Ptr8) are mutual best-hits, suggesting Tar2 represents ray parenchyma cells. From the scRNA-seq results of the three species in eudicots, we identified many marker genes from the DEG lists with relatively exclusive gene expression of different cell clusters (Extended Data Figs. 18, 19, Supplementary Tables 5, 6). Notably, all cell clusters in *T. aralioides* show very low correlation to libriform fibers (Ptr7) (Fig. 2f), which implies that the fusiform lineage in *T. aralioides* only contains one path toward tracheid/vessel characteristics. Namely, tracheids and vessel elements share a similar developmental path. Tracheids have both water transportation and mechanical support functions, and they became the specialized functions of vessel elements and libriform fibers, respectively^2^. Based on the transcriptomic analyses, the water transportation function, instead of mechanical support, is the main feature maintained by both tracheid- and vessel-bearing plants.

### Common and distinct developmental trajectories

The cell lineages in the four woody species were further compared by integrating their scRNA-seq data using the Seurat canonical correlation analysis pipeline^46^ for two-species graph-based cell clustering using orthologous genes (Fig. 3). Comparable to single-species clustering (Fig. 3a (i)–(iv)), *P. trichocarpa* and *E. grandis* two-species clustering (Fig. 3a (v), (vi)) shows highly overlapped (Fig. 3a (vii)) and similar fusiform and ray cell lineages (Fig. 3a (viii)). Egr6 was grouped with the ray organizer (Ptr3) in the two-species clustering (Fig. 3a (v)–(viii)), and was also the best-hit cluster of Ptr3 (Fig. 2e). We thus can also observe two cell lineages in the SDX of *E. grandis*, which are highly similar to that of *P. trichocarpa* differentiated from two types of organizer cells (Fig. 3a).

**Fig. 3.**
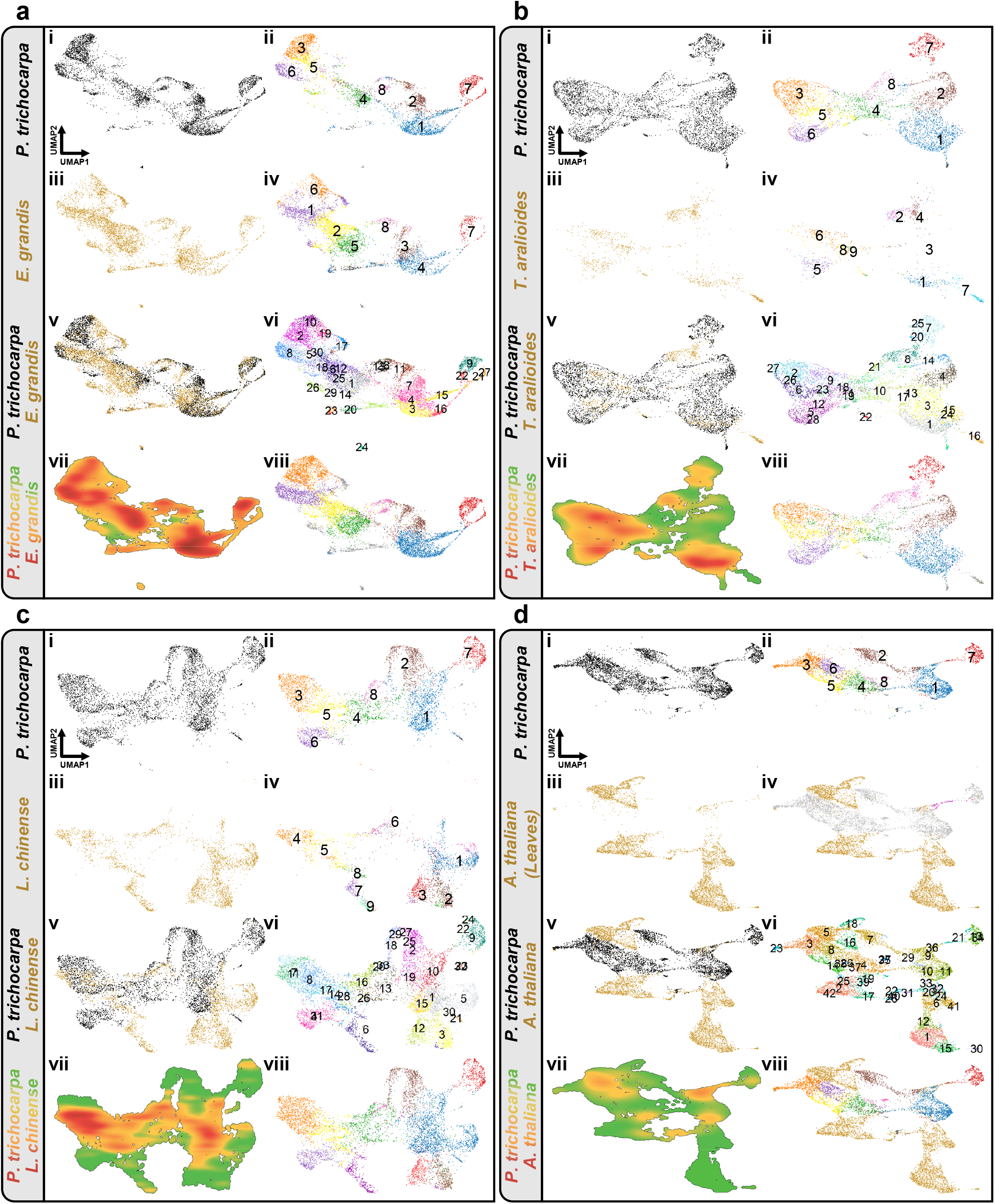
Two-species clustering and visualization of scRNA-seq data between *P. trichocarpa* and *E. grandis*, *T*. *aralioides*, *L. chinense* or *A. thaliana*. a–c, Two-species clustering of SDX single cells in *P. trichocarpa* and *E. grandis*. (**a**), *T. aralioides* (**b**) or *L. chinense* (**c**). (i)–(iv), Single-species unsupervised K-means clustering. (v)–(viii), Two-species graph-based cell clustering using orthologous genes. (i), (iii), (v), Black dots are SDX cells from *P*. *trichocarpa* and gold dots are cells from *E*. *grandis*, *T*. *aralioides*, *L. chinense*. (ii), (iv), The colors of cell clusters for each species are based on their single-species cell clustering results. (vi), The cell clusters in two-species clustering. (vii), The cell density of the two-species clustering. Green to orange to red colors indicate the density from low to high. (viii), The colors of two-species clustering are derived from that of single-species clustering (see Methods). **d**, Two-species clustering of SDX cells in *P. trichocarpa* and leaf cells in *Arabidopsis thaliana*. (i)–(iii), Single-species unsupervised K-means clustering. (iv)–(viii), Two-species graph-based cell clustering using orthologous genes. (i), (v), Black dots are SDX cells from *P*. *trichocarpa*. (iii), (iv), (v), (viii), Gold dots are cells from *A. thaliana*. (iv), Grey dots represent the SDX cells from *P. trichocarpa*, and the xylem cells identified in previous *Arabidopsis* studies are in magenta. (ii), (viii), The colors of cell clusters are based on the single-species cell clustering results. (vi), The cell clusters in two-species clustering. (vii), The cell density of the two-species clustering.

Two-species clustering of *P. trichocarpa* and *T. aralioides* also supports the conclusion from pairwise correlation analysis (Fig. 3b). Almost no cells from *T. aralioides* are grouped with libriform fibers (Ptr7) of *P. trichocarpa*, and Tar1 and Tar3 overlap with *P. trichocarpa* vessel elements (Ptr1) in UMAP. The fusiform lineage in *T. aralioides* has fusiform organizer (Tar5), fusiform early precursor (Tar9), fusiform intermediate precursor cells (Tar3) and fusiform late precursor/tracheid cells (Tar1) (Fig. 3b (v)–(viii)). The ray lineage started from the ray organizer (Tar6) to ray precursor (Tar8) then to ray parenchyma cells (Tar2) (Fig. 3b (v)–(viii)). Based on the similar developmental paths among the three woody eudicots, we could identify many distinct modules of orthologous groups of genes with temporal expression patterns along the pseudotime of each cell lineage in two-species analyses (Extended Data Figs. 20, 21, Supplementary Tables 7, 8).

In the two-species clustering of *P. trichocarpa* and *L. chinense*, we found that the ray lineage is highly similar, but the fusiform lineage in *L. chinense* exhibits a distinct path from that in *P. trichocarpa* (Fig. 3c). Cell clusters corresponding to fusiform early (Ptr4/Lch8), intermediate (Ptr2/Lch2) and late precursors/vessel elements (Ptr1/Lch1) barely overlap between *P. trichocarpa* and *L. chinense* (Fig. 3c (v)–(viii)). Despite their low transcriptomic correlation (Fig. 2g), fusiform organizer cells (Lch7) and libriform fibers (Lch3) of *L. chinense* could be identified through plotting the marker genes from *P. trichocarpa* (Extended Data Fig. 22), suggesting that some key genes involved in fusiform cell lineage development are shared by the two distantly related angiosperms.

Overlapping cell populations between *P*. *trichocarpa* and each of the other three plants provide important insights into similarities and variation in developmental trajectories. To test whether such overlap could be caused by over-correction for dimensionality reduction in two-species analysis, we conducted the same procedure for *P. trichocarpa* SDX cells with each of the previously reported scRNA-seq data from *Arabidopsis* leaves (Fig. 3d), *Arabidopsis* roots (Extended Data Fig. 23a) and rice (*Oryza sativa*) roots (Extended Data Fig. 23b), where a sizable number of xylem or vascular cells were present (Extended Data Fig. 24). Most of the SDX cells from *P. trichocarpa* could not be grouped with the root and leaf cells from *Arabidopsis* and rice (Fig. 3d, Extended Data Fig. 23a, b), and the densities of overlapping cells between *P. trichocarpa* and the other three woody species are much higher than those between *P. trichocarpa* and *Arabidopsis* or rice (Fig. 3a (vii), b (vii), c (vii), d (vii), Extended Data Fig. 23a (vii), b (vii)). All of the xylem marker genes identified in previous scRNA-seq studies could be detected in our SDX scRNA-seq dataset (Extended Data Fig. 25). The distribution of the root and leaf xylem cells (magenta, Fig. 3d (iv), Extended Data Fig. 23a (iv), b (iv)) only barely overlaps with that of the *P*. *trichocarpa* SDX fusiform late precursors/vessel elements (Ptr1), libriform fibers (Ptr7), and the transition area of these two clusters. In addition, almost no *Arabidopsis* xylem cells are co-localized with the terminal differentiated stage libriform fiber cluster (Fig. 3d (iv), Extended Data Fig. 23a (iv)), which supports the notion that the fiber cells in *Arabidopsis* only have an incomplete maturation process^47^. The *Arabidopsis* xylem cells also do not overlap with *P*. *trichocarpa* ray parenchyma cells (Ptr8) (Fig. 3d (iv), Extended Data Fig. 23a (iv)), consistent to previous finding that ray parenchyma cells cannot be observed in *Arabidopsis* under normal growing conditions^48–50^. The relatively limited xylem cells from previous studies of *Arabidopsis*, which has incomplete xylem development and cell types, provide only a peek into the tip of the iceberg of xylem transcriptomic variation. Overall our results demonstrate conserved and overlapping developmental paths of SDX across divergent woody angiosperms, whereas the developmental trajectories of SDX in the rosid *P*. *trichocarpa* are clearly distinct from those of root or leaf xylem in another rosid *Arabidopsis*.

The conclusion from single- (Figs. 1, 2) and two-species analyses (Fig. 3) is further supported by four-species combined analyses (Fig. 4). All four angiosperms have nearly identical ray cell lineages (Fig. 4f–i). The three eudicots share similar fusiform cell lineages (red lines, Fig. 4j, k, m), except that *T. aralioides* does not have libriform fibers (Fig. 4l). *L. chinense* in magnoliids has a distinct fusiform cell lineage due to its different fusiform organizer cells and libriform fibers (Fig. 4m).

**Fig. 4.**
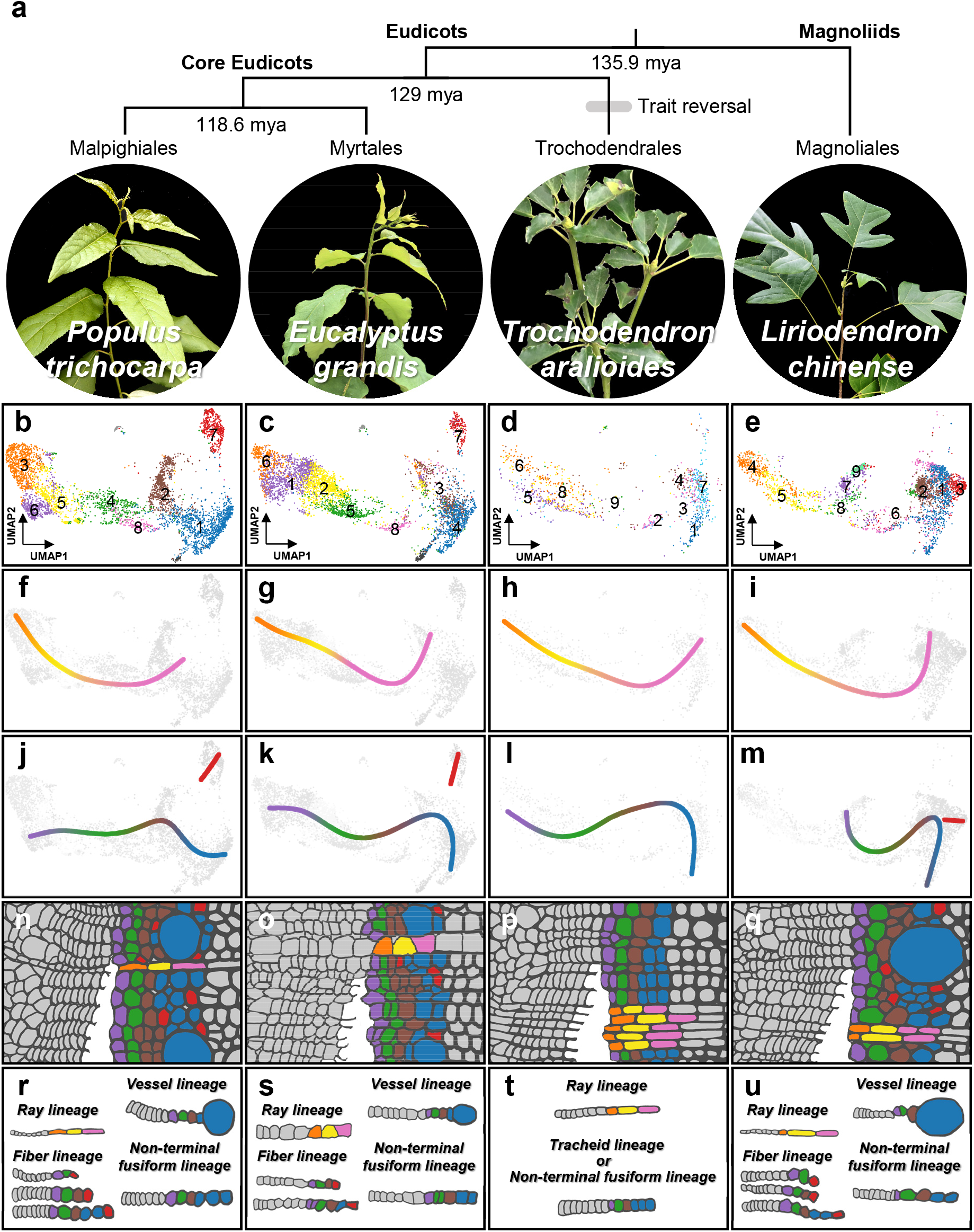
Cell lineages in SDX development in four woody angiosperms. **a**, Phylogenetic relationships and estimated divergence times^5^ of *P. trichocarpa*, *E. grandis* (both core eudicots), *T. aralioides* (basal eudicots) and *L. chinense* (magnoliids). **b**–**e**, Combined four-species analyses and two-dimensional (2D) visualization of SDX scRNA-seq data. Individual cells are colored as in Figs. 1, 2. **f**–**m**, The ray (**f**–**i**) and fusiform (**j**–**m**) lineages in the four species. **n**–**u**, Schematics of different cell lineages across the four species.

Using lcmRNA-seq and scRNA-seq of multiple species, this study provides crucial insights into the formation and developmental dynamics of SDX. In particular, we discovered that (1) the radial (ray parenchyma cells) and axial (libriform fibers and tracheids/vessel elements) systems in SDX develop through two cell lineages from ray and fusiform organizer cells to precursor cells then to terminally differentiated cells (Fig. 4n–u); (2) The transcriptomic profile of tracheids is most similar to that of vessel elements, suggesting a more dominant role of water transportation in fusiform development than mechanical support; (3) In contrast to highly conserved ray lineages, fusiform lineages have variable features across core eudicots, basal eudicots, and magnoliids (Fig. 4). SDX constitutes the largest biomass on Earth as the essential tissue for two major functions in vascular plants. The mechanical support allows vascular plants to dominate current terrestrial ecosystem by towering over their bryophyte relatives. Water transportation delivers critical elements to whole plant bodies. By integrating high-resolution single-cell analyses across multiple species with evolutionary divergence of over a hundred million years, we provide a comprehensive view of how cells form and vary in SDX. Further studies of the entire plant kingdom could shed light on the origin of vascular development.

## Methods

### Plant materials and sampling

*P. trichocarpa* plants were grown in a walk-in growth chamber under 16-hour light/8-hour dark conditions at 24–26°C. Tree stems were collected from 8-month-old plants. For protoplast isolation, stems below the 12^th^ internode were used. For LCM, the 8^th^ internode of the tree stems was harvested. *E. grandis* plants were grown in a walk-in greenhouse for 8 months with sunlight and temperature around 20–30°C. Tree stems below the 18^th^ internode were collected for protoplast isolation. The branches from wild-grown *T. aralioides* in Hutian Elementary School (Yangmingshan National Park, Taiwan) were collected for protoplast isolation. Around 1-year-old plants of *L. chinense* were purchased from Nanping Senke Seedlings Co., Ltd. (Fujian, China), and the stems below around the 10^th^ internode were used for protoplast isolation.

### SDX protoplast isolation

The SDX protoplast isolation was carried out mainly based on our previously established protocol^51–53^. Tree stems were cut into 8-centimeter segments. Debarked stem segments were dipped into freshly prepared enzyme solution (20 mM MES, pH 5.7, 0.25 M mannitol, 20 mM KCl, 1.5% wt/vol cellulase R10, 0.4% wt/vol Macerozyme R10, 10 mM CaCl_2_ and 0.1% BSA) for ∼3 hours at room temperature. Enzyme-digested stem segments were transferred to MMG solution (4 mM MES, pH 5.7, 0.25 M mannitol and 15 mM MgCl_2_, room temperature) to release SDX protoplasts. The protoplasts were filtered with 70-μm nylon mesh, pelleted at 900 g centrifugation for 3 minutes at room temperature, and resuspended in MMG solution.

### 10x Chromium scRNA-seq

Single-cell sequencing libraries of one biological replicate for *P. trichocarpa* and *L. chinense* SDX protoplasts were generated using Chromium Single Cell 3’ Library and Gel Bead Kit v2 (10x Genomics) according to user manual. The scRNA-seq libraries of *P. trichocarpa* and *L. chinense* were sequenced using Illumina HiSeq4000 (Genomics Co., Ltd) and NovaSeq (Novogene Co., Ltd), respectively. The sequenced data of *P. trichocarpa* and *L. chinense* were individually processed using Cell Ranger (10x Genomics; v5.0.1) with the commands “cellranger mkref” and “cellranger count”. The Cell Ranger-compatible references were built with the current genome sequences and annotations (see below) for the following read alignment and the quantification. The single-cell transcript abundance matrices were generated and named as “filtered_feature_bc_matrix.h5” by Cell Ranger. The transcript abundance of each gene was calculated based on the number of unique molecular identifier (UMI)-tagged transcripts as “UMI counts”. A total of 4,705 and 2,977 transcriptomes were obtained for individual SDX cells in *P. trichocarpa* and *L. chinense*, respectively, each with UMI counts more than 500 (Extended Data Fig. 26). In *P. trichocarpa*, total 27,102 genes were detected among the transcriptomic dataset, which represents 78.1% of the whole annotated genes (34,699 genes) (Extended Data Fig. 26). In *L. chinense*, 17,926 genes were detected, representing 50.8% of the whole annotated genes (35,269 genes).

### Cell sorting and plate-based scRNA-seq

The massively parallel single cell RNA-seq2.0 (MARS-seq2.0)^54^ protocol was adapted for sequencing *E. grandis* and *T. aralioides* SDX protoplasts using one biological replicate for each species. 384-well plates were prepared using Beckman Biomek NXR Liquid Handling Workstation (Beckman, USA), with 2 µL lysis buffer (0.1% Triton X-100, 0.2U/µL RNAsin Plus and 1 nM reverse transcription (RT) primers with barcodes) in each well. *E. grandis* and *T. aralioides* SDX protoplasts were stained with a working concentration of 0.008 µg/µL fluorescein diacetate (Sigma-Aldrich, USA). The samples were analyzed and sorted by a Beckman Coulter MoFlo XDP (Beckman, USA) or BD FACSMelody™ cell sorter (BD, USA) with 488-nm laser excitation and a 100-µm nozzle. Cells were identified based on forward scatter (FSC), side scatter (SSC), chlorophyll fluorescence (663–737 nm) and fluorescein fluorescence (475–650 nm). A single cell was isolated under the “single cell mode” into individual wells of the prepared 384-well plates. After sorting, the plates were immediately frozen on dry ice and stored at −80°C until processing. The plates were thawed at room temperature, heated to 95°C for 3 minutes, and cooled on an iced metal block. The RT reagent mixture, including 1:80,000-diluted ERCC spike-in RNA, was dispensed into each well using Mantis v3.2 (Formulatrix, Massachusetts, USA). After RT and Exonuclease I treatment, all wells in each plate were pooled together through 3-minute centrifugation at 1,800 rpm (Heraeus Multifuge X3R with a TX-750 swinging bucket rotor, Thermofisher, USA). Further processing of the plate pools followed the MARS-seq2.0 protocol, including second-strand synthesis, *in vitro* transcription, DNase treatment, RNA fragmentation, ligation and plate barcoding and second RT. The quality of the libraries was assessed by quantitative PCR using QuantStudio 12K Flex Real-Time PCR System (Applied Biosystems, USA). The cDNA pools were amplified by a final PCR with the introduction of Illumina I7 indices. Libraries were paired-end sequenced (R1:130, R2:20) on Illumina NextSeq 500 (Genomics Core, Institute of Molecular Biology, Academia Sinica, Taiwan). Sequencing data was analyzed using the MARS-seq2.0 mapping and demultiplexing pipeline^54^. In *E. grandis*, total 22,217 genes were detected among the transcriptomic dataset, which represents 61.1% of the whole annotated genes (36,349 genes) (Extended Data Fig. 26). In *T. aralioides*, 23,916 genes were detected, representing 67.7% of the whole annotated genes (35,328 genes).

### Dimensionality reduction and cell clustering

For *P. trichocarpa* and *L. chinense*, the dimensionality reduction and cell clustering were performed by Cell Ranger (10x Genomics; v5.0.1) using the single-cell transcript abundance matrices with the command “cellranger reanalysis”. For *E. grandis* and *T. aralioides*, the results from MARS-seq2.0 were converted into the single-cell transcript abundance matrices as the input files for Cell Ranger to exert the subsequent command “cellranger reanalysis” for dimensionality reduction and cell clustering. The principal component analysis was performed, and the top 10 principal components were used to conduct UMAP and unsupervised K-means clustering with the parameters “umap_n_neighbors = 50” and “umap_min_dist = 0.5”. Differential expression analysis was performed using two methods. Negative binomial exact test (two-sided), the hypothesis test in default, based on the sSeq method^55^, was applied to identify the DEGs in each cluster comparing to other clusters. When the counts of transcript abundance in both comparing groups were greater than 900, the hypothesis test would be switched to asymptotic beta test (two-sided) adapted from the implementation in edgeR^56^. The *P* values from the multiple comparisons were corrected by Benjamini-Hochberg method (BH method)^57^ to control the false discovery rate (FDR) as 0.05. DEGs of each cell cluster were obtained using the output table “differential_expression.csv” in the folder “kmeans_X_clusters” generated by Cell Ranger. The “X” in “kmeans_X_clusters” was corresponding to the cluster number in each species. Only the clusters with more than or equal to 5 cells would be considered as cell clusters. The cutoff of DEGs was set as adjusted *P* value < 0.05 and the absolute value of log_2_(fold change) ≥ 1. The UMI counts of each cell were divided by the total UMI counts and multiplied by a scale factor (1,000) to obtain the normalized UMI counts. The log_2_-transformation was applied to the normalized UMI counts to plot the gene transcript abundance.

### Laser capture microdissection and RNA sequencing (lcmRNA-seq)

The 8^th^ internodes from six *P. trichocarpa* stems were cut into 5- or 10-mm segments, frozen by liquid nitrogen immediately, and stored at −80°C. The 5-mm segments were separated into four quarters at −20°C in the chamber of a cryostat. The segments were stuck on cryostat chucks using Tissue-Tek O.C.T. Compound (Sakura Finetek, USA), and sliced into 16-μm thick sections using cryostat (Leica CM1900) at −20°C. The cryo-sections were placed onto the PET membrane of metal-frame slides (Leica, USA), and dehydrated using 99.5% ethanol. Around 1,600,000 µm^2^ area from cryo-sections (cross or tangential) of two *P. trichocarpa* plants was collected using laser microdissection systems (Leica LMD7000), and evenly pooled as one biological replicate. A total of three biological replicates were collected for each cell type. The dissected cells were collected within 30 minutes in 500-μL eppendorf caps loaded with 50 µL RLT buffer (QIAGEN, USA) and 1% beta-mercaptoethanol (Sigma-Aldrich, USA), frozen by liquid nitrogen, and stored at −80°C. Total RNA from dissected cells was extracted using RNeasy Plant Mini Kit (QIAGEN, USA) according to the manufacturer instruction. The mRNA of each cell type was amplified using Arcturus® RiboAmp® HS PLUS RNA Amplification Kit (Applied Biosystems, USA) following the user guide. The amplified mRNA of each sample was used to construct sequencing libraries using NEBNext® Ultra™ II Directional RNA Library Prep Kit (New England Biolabs, USA) according to the instruction manual. The RNA-seq libraries were sequenced on Illumina NextSeq 500 platform (Genomics Core, Institute of Molecular Biology, Academia Sinica, Taiwan) to obtain the reads with paired-end 75-bp length.

### Microscopic imaging

The transverse and tangential sections were observed under Olympus BX53 Upright Microscope in bright field. The sections for LCM were examined using the embedded camera function in laser microdissection systems (Leica LMD7000).

### lcmRNA-seq data analysis

The sequencing raw reads were preprocessed with fastp^58^ (v0.20.1) using five arguments “--detect_adapter_for_pe, --correction, --cut_front, --cut_tail and -- disable_trim_poly_g”. The adapter sequences and low-quality sequences (mean quality value < 20) at the both 5’ and 3’ ends of each read were removed. The reads with any of the following features were discarded: (1) shorter than 15 bps, (2) containing more than five N bases and (3) including more than 40% of bases with low quality (quality value < 15). The filtered reads were mapped to the *P. trichocarpa* reference genome with HISAT2^59, 60^ (v2.2.1) using four arguments “--max-intronlen 17787, --secondary, --fr and --rna-strandness”. The parameter of the argument “--max-intronlen 17787” as maximum intron length was obtained from the current *P. trichocarpa* genome structural annotation (see below). The transcriptome quantification was performed using StringTie^61^ (v2.1.3b) with the current annotation to obtain the raw read counts and transcripts per million (TPM) of each gene. The differential expression analysis was implemented with the negative-binomial-based test (two-sided) in DESeq2^62^ (v1.28.1) using the raw read counts. The FDR was controlled with BH method^57^, and the significance level was set as 0.05. DEGs of each cell type were identified under the cutoff set as the absolute value of log_2_(fold change) ≥ 1 compared with the other cell types (Supplementary Table 9). For example, the DEGs of libriform fibers were the genes with the absolute value of log_2_(fold change) ≥ 1 both in libriform fibers/vessel elements and libriform fibers/ray parenchyma cells.

### Correlation analysis between scRNA-seq and lcmRNA-seq data

The Pearson’s correlation coefficients were calculated between the transcriptome of each cell from scRNA-seq and each cell type from lcmRNA-seq data. UMI counts and average TPM were used to represent the transcript abundance of the transcriptome data of each cell in scRNA-seq and each cell type in lcmRNA-seq, respectively.

### Transcript abundance quantification of SDX

The RNA-seq raw reads from three biological replicates of SDX in *P. trichocarpa* were downloaded from NCBI GEO database using the accession number GSE81077^63^. The RNA-seq data was processed using the same pipeline as lcmRNA-seq data analysis mentioned above to obtain TPM of each gene in SDX.

### Identification of homologous genes

We used sequence similarities to identify homologous protein-coding genes encoded in genomes of the four woody angiosperms and 10 other species with a balanced coverage of the plant diversity, including *Physcomitrium patens*, *Marchantia polymorpha*, *Selaginella moellendorffii*, *Pinus taeda*, *Gnetum montanum*, *Amborella trichopoda*, *Oryza sativa*, *Arabidopsis thaliana*, *Coffea canephora* and *Solanum lycopersicum*. An all-against-all BLASTP search^64, 65^ (v2.6.0) was conducted for the amino acid sequences with an e-value cutoff of 10^-15^. OrthoMCL^66^ (v1.3) was used to cluster the protein-coding genes based on their BLASTP similarities into orthologous groups using the Markov Cluster Algorithm^67^ with an inflation of 1.5 (Supplementary Table 10). For the cross-species analyses, the ortholog transcript abundance was calculated as the total gene transcript abundance within each orthologous group. The ortholog transcript abundance was normalized with the total ortholog transcript abundance in each cell, and then multiplied by a scale factor (1,000). The log_2_-transformation was applied to the normalized ortholog transcript abundance to plot the ortholog transcript abundance.

### Pairwise correlation analysis among cell clusters

For *P. trichocarpa*, the Pearson’s correlation coefficients were calculated between each pair of major cell clusters (Ptr1–Ptr8) using the average normalized UMI counts within each cell cluster. For two-species comparisons, the average normalized ortholog transcript abundance was calculated within each major cell cluster (except Egr9, Egr10, Tar10 and Lch10) for generating the Pearson’s correlation coefficients between each pair of cross-species cell clusters.

### MetaCell analysis of *P*. *trichocarpa* SDX cells

As an alternative to unsupervised K-means clustering and UMAP visualization, the MetaCell package^41^ (v0.3.41) was used to group the *P*. *trichocarpa* SDX cells and project them onto a 2D plot. A total of 4,278 variable genes were selected as marker genes (T_tot = 20, T_top3 = 1, T_szcor = −0.01, T_vm = 0.2, and T_niche = 0.05) for constructing k-nearest neighbor graphs with K = 40, followed by coclustering with bootstrapping based on 1000 iterations of resampling 75% of the cells and an approximated target minimum metacell size of 50. Unbalanced edges were filtered with K = 40 and alpha = 3. The individual cells were then plotted in a 2D projection (Extended Data Fig. 12).

### Two-species or four-species scRNA-seq data integration for four woody plants

Seurat pipeline (v4.0.3) was used for the integration for the scRNA-seq results from multiple species^46, 68^. Only the cells with more than 200 detected orthologs and the orthologs detected in more than 3 cells were included in the analysis. The ortholog transcript abundance in the filtered data was normalized according to Seurat guideline. Top 2,000 highly variable orthologs were identified in each species, and used as the input of canonical correlation analysis (CCA). The correlated gene modules were identified and presented as the cross-species cells embedding in a shared low-dimensional space. L2-normalization was then performed on the cell embedding to reduce the global different between datasets. The anchors, defined as the mutual nearest neighbors between datasets, were identified with K-nearest neighbors (K = 5) using 2000 highly variable orthologs among two or four species. After anchor scoring and anchor weighting, the anchors were then used for data integration. UMAP and cell clustering were performed based on shared nearest neighbor modularity optimization. The determination for the cell clusters (color) in two-species clustering (Fig. 3a (viii), b (viii), c (viii)) was mainly based on the majority cell clusters (color) from single-species clustering results. Take a cell cluster in two-species clustering for *P. trichocarpa* and *E. grandis* as an example, if this cell cluster contains 100 cells, 50 from *P. trichocarpa* (48 in red cluster and 2 in brown cluster) and 50 from *E. grandis* (30 in red cluster and 20 in brown cluster), then all 100 cells would be defined as red cell cluster.

### Re-analysis of *Arabidopsis* and rice scRNA-seq data

For *Arabidopsis* leaves^22^, the scRNA-seq data of one biological replicate (genotype ATML1p::YFP-RCI2A) obtained by 10x Genomics v2 was downloaded from NCBI GEO database using the accession number GSE167135 as single-cell transcript abundance matrices in one file (filtered_gene_bc_matrices-_h5.h5). This h5 file was analyzed with the command “cellranger reanalysis” in Cell Ranger (10x Genomics; v5.0.1) to obtain the UMAP and unsupervised K-means clustering results (Extended Data Fig. 27a). According to Lopez-Anido et al.^22^, the xylem cells in *Arabidopsis* leaves can be identified using the gene expression of *AT3G21270* (Extended Data Fig. 27a). For *Arabidopsis* roots^13^, the scRNA-seq data with three biological replicates of wild type was downloaded from NCBI GEO database under the accession number GSE123013 as single-cell transcript abundance matrices composed of three separated files (barcodes.tsv, genes.tsv and matrix.mtx). The three biological replicates of the wild-type cell transcriptomes were integrated and processed (Extended Data Fig. 27b) by Seurat as mentioned above. According to Ryu et al.^13^, the xylem cells in *Arabidopsis* roots can be identified using the gene expression of *BHLH30* (*AT1G68810*) (Extended Data Fig. 27b). For rice roots^21^, the scRNA-seq raw reads from two biological replicates of wild type were downloaded from NCBI SRA database under the accession number PRJNA706435. Transcriptome quantification, followed by the UMAP and unsupervised K-means clustering, was performed by Cell Ranger (10x Genomics; v5.0.1) using the command “cellranger mkref” and “cellranger count” (Extended Data Fig. 27c). According to Zhang et al.^21^, the xylem cells in rice roots can be identified using the gene expression of *Os01g0750300* (Extended Data Fig. 27c). Each re-analyzed scRNA-seq dataset of *Arabidopsis* roots and leaves and rice roots were integrated with the SDX scRNA-seq results of *P. trichocarpa* by Seurat using the pipeline mentioned above.

### Cell density of overlapped cells

The overlapping degree among the cells from Seurat integrating results was visualized based on the distribution of intra-species minimal spanning tree (MST) derived from previous studies^69, 70^. The MST was first constructed using the integrated UMAP (Extended Data Fig. 28a, b). The subgraphs were obtained by removing the cross-species edges (Extended Data Fig. 28c–f). The center nodes for each subgraph, defined as the node with the highest closeness centrality, were determined (Extended Data Fig. 28g, h). The MST construction, graph determination and closeness centrality calculation were performed using igraph^71^ (v1.2.6). For the single-node subgraph, such only node in the subgraph was directly defined as the center node (Extended Data Fig. 28g, h). The two-dimensional kernel density estimation was performed using MASS^72^ (v7.3-54) based on the distribution of center nodes of each species to obtain the subgraph center density map (Extended Data Fig. 28i, j). The density maps were then weighted by the number of center nodes and the proportion of cells of each species, and the weighted density maps were concatenated into a single density map. The density map was then normalized by the total density values of *P. trichocarpa* cells.

### Pseudotime trajectory analysis and lineage curve construction

The pseudotime analysis was performed by Slingshot^73^ (v2.0.0) based on the integrated UMAP from Seurat. From our defined cell lineages as tracheid-, vessel element-, libriform fiber- and ray parenchyma-lineages, the pseudotime of each cell in our defined cell lineages was obtained, and the cells were ordered based on their corresponding pseudotime. The window size was set as 21 cells and slid one cell at a time to calculate the moving average from the ortholog transcript abundance (UMI) in each cell. For example, one orthologous group in the first 21 cells with minimal pseudotime in a certain cell lineage would generate an average of transcript abundance, and such orthologous group in the 2^nd^ to 22^th^ cells would generate another average. The union of all of these averages is named as moving average. The differentially expressed orthologous groups (DEOs) were identified using the command “FindAllMarkers” in Seurat with MAST^74^ (v1.18.0), which was based on the hurdle model tailored to scRNA-seq analysis. Two-sided tests were performed, and the output *P* values were adjusted based on Bonferroni correction. Under the criteria of adjusted *P* value < 0.05, DEOs were obtained as the orthologous groups differentially expressed in any one of the integrated cell clusters from two species. Each DEO in different cell lineages was plotted using their corresponding moving averages. The DEOs from each cell cluster belong to different modules, yielding total eight modules from A to H. More than 95% of the cells in each cell cluster from every species were used to construct the smooth lineage curves basically passing the centers of each cell cluster according to the pre-defined lineages using Slingshot^73^.

### Data source of genome sequences, genome annotations and peptide sequences

We downloaded the genome sequences, genome annotations or peptide sequences of the 14 species from the following databases: *Populus trichocarpa* (v4.1; Phytozome v13, https://phytozome-next.jgi.doe.gov/), *Eucalyptus grandis* (v2.0; Phytozome v13), *Trochodendron aralioides* (released on October 10^th^ 2019; GigaDB, http://gigadb.org/), *Liriodendron chinense* (generated on January 15^th^ 2019; Hardwood Genomics Project, https://hardwoodgenomics.org/), *Amborella trichopoda* (Ensembl Genomes, release 49, http://ftp.ensemblgenomes.org/pub/plants/release-49/), *Arabidopsis thaliana* (Araport11; The Arabidopsis Information Resource, https://www.arabidopsis.org/), *Coffea canephora* (Coffee Genome Hub, https://coffee-genome.org/), *Gnetum montanum* (Dryad Digital Repository, https://datadryad.org/), *Marchantia polymorpha* (MpTak1v5.1; MarpolBase, https://marchantia.info/), *Oryza sativa* (Ensembl Genomes, release 49), *Physcomitrium patens* (v3.3; Phytozome v13), *Pinus taeda* (v2.01; TreeGenes, https://treegenesdb.org/), *Selaginella moellendorffii* (v1.0; Phytozome v13), *Solanum lycopersicum* (ITAG4.0; Sol Genomics Network, https://solgenomics.net/).

## Acknowledgements

We acknowledge Technology Commons in College of Life Science and the Instrumentation Center sponsored by Ministry of Science and Technology, National Taiwan University for technical assistance. We are grateful to Hutian Elementary School for providing the plant materials of *T. aralioides*. Y.J.L. was supported by Young Scholar Fellowship Columbus Program, Ministry of Science and Technology of Taiwan (MOST) (107-2636-B-002-003, 108-2636-B-002-003, and 109-2636-B-002-003). C.K. was supported by Academia Sinica Career Development Award (AS-CDA-110-L01) and a MOST research grant (108-2311-B-001-040-MY3). J.C.S. was supported by Young Scholar Fellowship Einstein Program, MOST (107-2636-B-010-002, 108-2636-B-010-002, and 109-2636-B-010-008). P.S. was supported by National Natural Science Foundation of China (Grant No. 31971626).

## Author information

These authors contributed equally: Chia-Chun Tung, Shang-Che Kuo, Chia-Ling Yang

## Contributions

Y.J.L., C.K. and Y.H.S. conceived the study. Y.J.L. carried out the protoplast isolation of *P. trichocarpa*. C.C.T., P.S. and Y.J.L. performed the protoplast isolation of *L. chinense*. C.C.T., S.C.K. and Y.J.L. performed the protoplast isolation of *E. grandis* and *T. aralioides*. C.C.T., C.L.Y. and C.K. conducted cell sorting of *E. grandis* and *T. aralioides*. C.L.Y. and C.K. carried out the plate-based scRNA-seq experiments of *E. grandis* and *T. aralioides*. C.E.H. contributed to the lcmRNA-seq experiments and microscopic imaging. S.C.K. and C.C.T. performed the lcmRNA-seq data analyses.

C.K. and C.L.Y. carried out the MARS-seq2.0 data analyses and identified the homologous genes. S.C.K. and C.K. performed single-species scRNA-seq data analyses. S.C.K., C.C.T. and Y.J.L. carried out the correlation analyses and cross-species scRNA-seq data analyses. Y.J.L., C.K., C.C.T and S.C.K. wrote the manuscript. C.C.T., S.C.K., Y.J.L., C.E.H. and C.K. prepared the figures. C.K., Y.J.L., C.C.T., S.C.K., C.L.Y., J.C.S. and J.H.Y. revised the manuscript. Y.J.L., C.K., J.C.S. and P.S. acquired the funding. Y.J.L. and C.K. supervised the study.

## Corresponding author

Correspondence to Chuan Ku or Ying-Chung Jimmy Lin

## Additional information

Supplementary Information is available for this paper.

## Extended data figures

**Extended Data Fig. 1.**
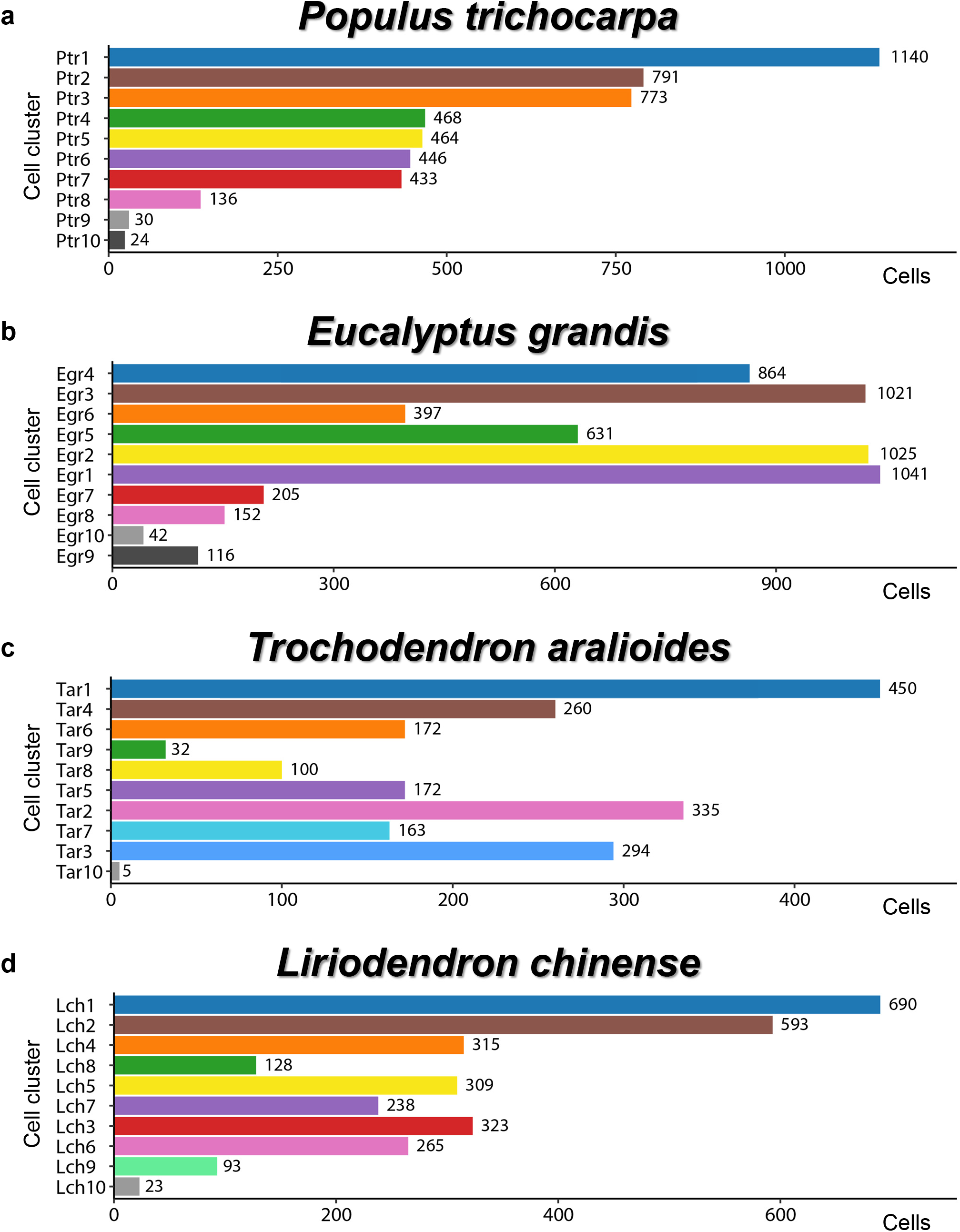
Cell numbers of SDX scRNA-seq cell clusters in *P. trichocarpa*, *E. grandis*, *T. aralioides* and *L. chinense*. **a**–**d,** SDX single-cell transcriptomes of *P. trichocarpa* (**a**), *E. grandis* (**b**), *T. aralioides* (**c**) and *L. chinense* (**d**) were grouped using unsupervised K-means clustering into 10 cell clusters in each species. The number of cells is shown for each cell cluster.

**Extended Data Fig. 2.**
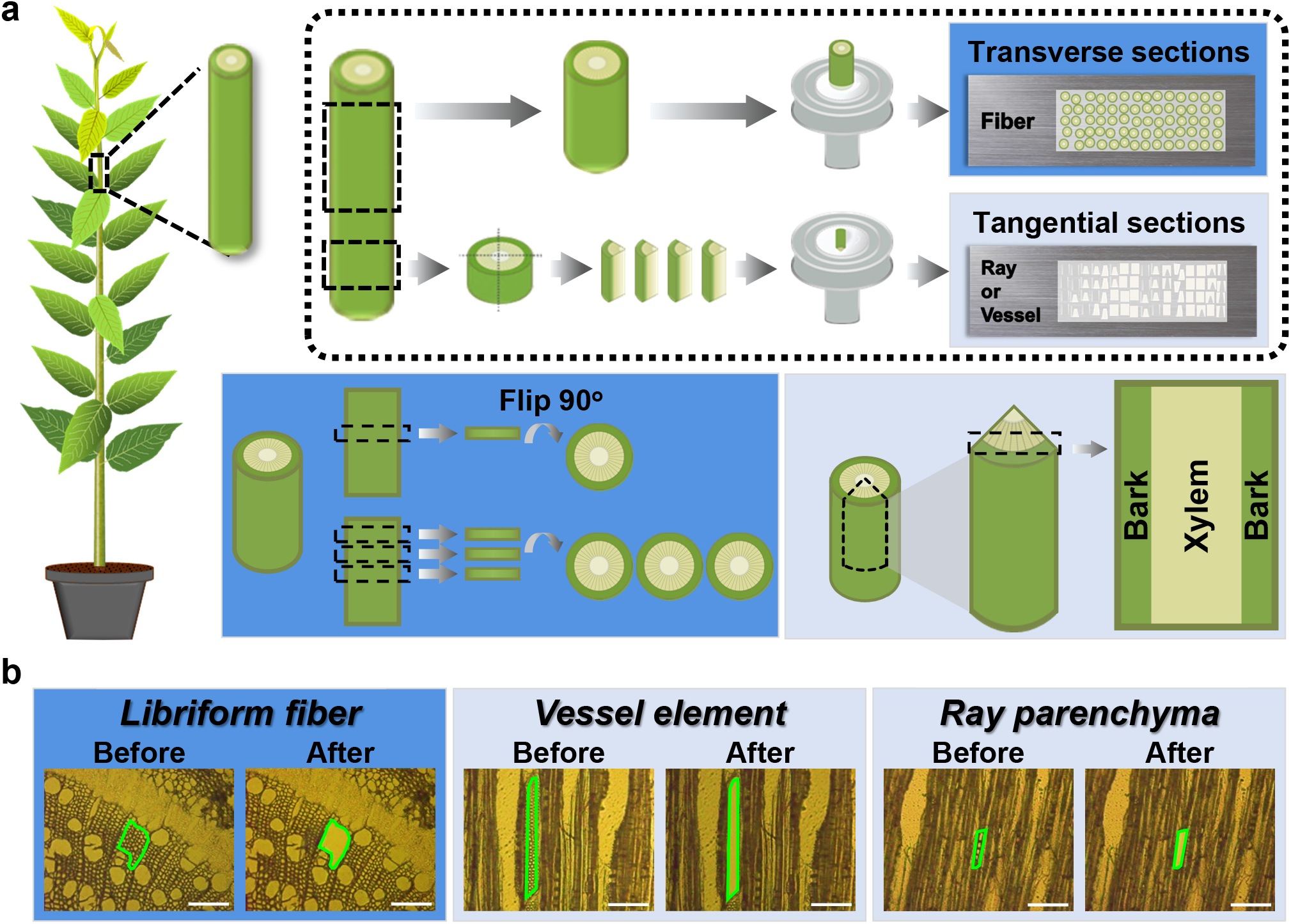
Workflow of laser capture microdissection for three xylem cell types in *P. trichocarpa*. **a**, Schematic of transverse and tangential sectioning of tree stems. A whole stem segment and a quarter stem segment are loaded on cryostat chucks for transverse and tangential sectioning, respectively. Sections are then placed on metal-frame slides with PET membrane for subsequent LCM cell type isolation. **b**, Real sections before and after laser cutting of libriform fibers, vessel elements and ray parenchyma cells. The area within the green circles represents the cutting area. Scale bars, 100 μm. **a**, **b**, Transverse and tangential sections are marked with dark-blue and light-blue backgrounds, respectively.

**Extended Data Fig. 3.**
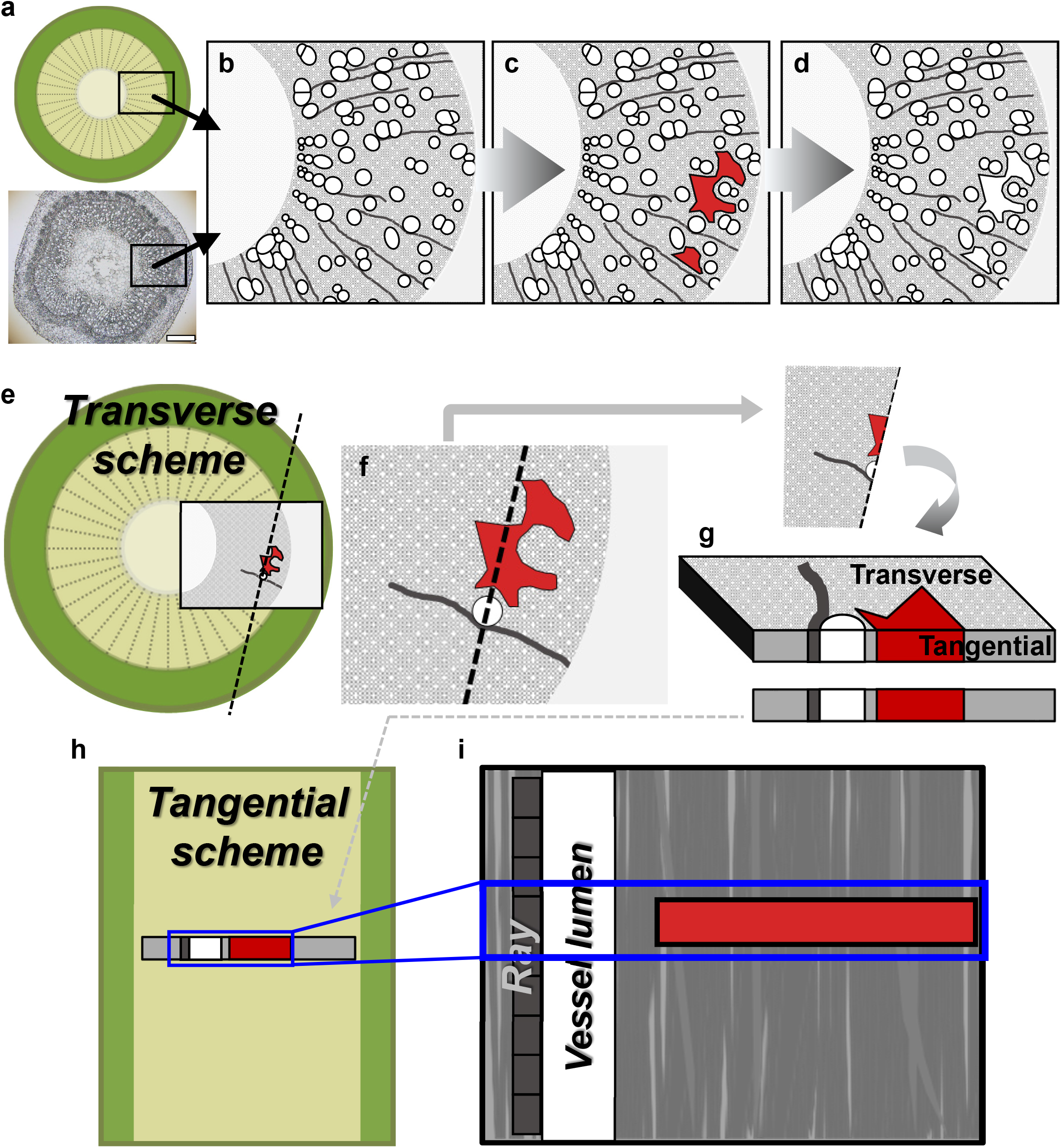
Libriform fiber collection in transverse and tangential views. **a**–**i**, The red area represents the libriform fibers collected using LCM. **a**–**d**, Libriform fiber collection from transverse sections, showing before cutting (**b**), area selection (**c**), and after cutting (**d**). Scale bar, 500 μm. **c**, **d**, The area switched from red to white, leaving an empty space after the libriform fibers were cut by laser. **e**–**i**, Libriform fibers in transverse and tangential views. **e**, A transverse section with an area highlighted (red color) for libriform fiber collection. **f**, A closeup of the highlighted area. **g**, Three-dimensional (3D) structure around the highlighted area. **h**, A tangential section with highlighted area. **i**, Corresponding panel of Fig. 1e.

**Extended Data Fig. 4.**
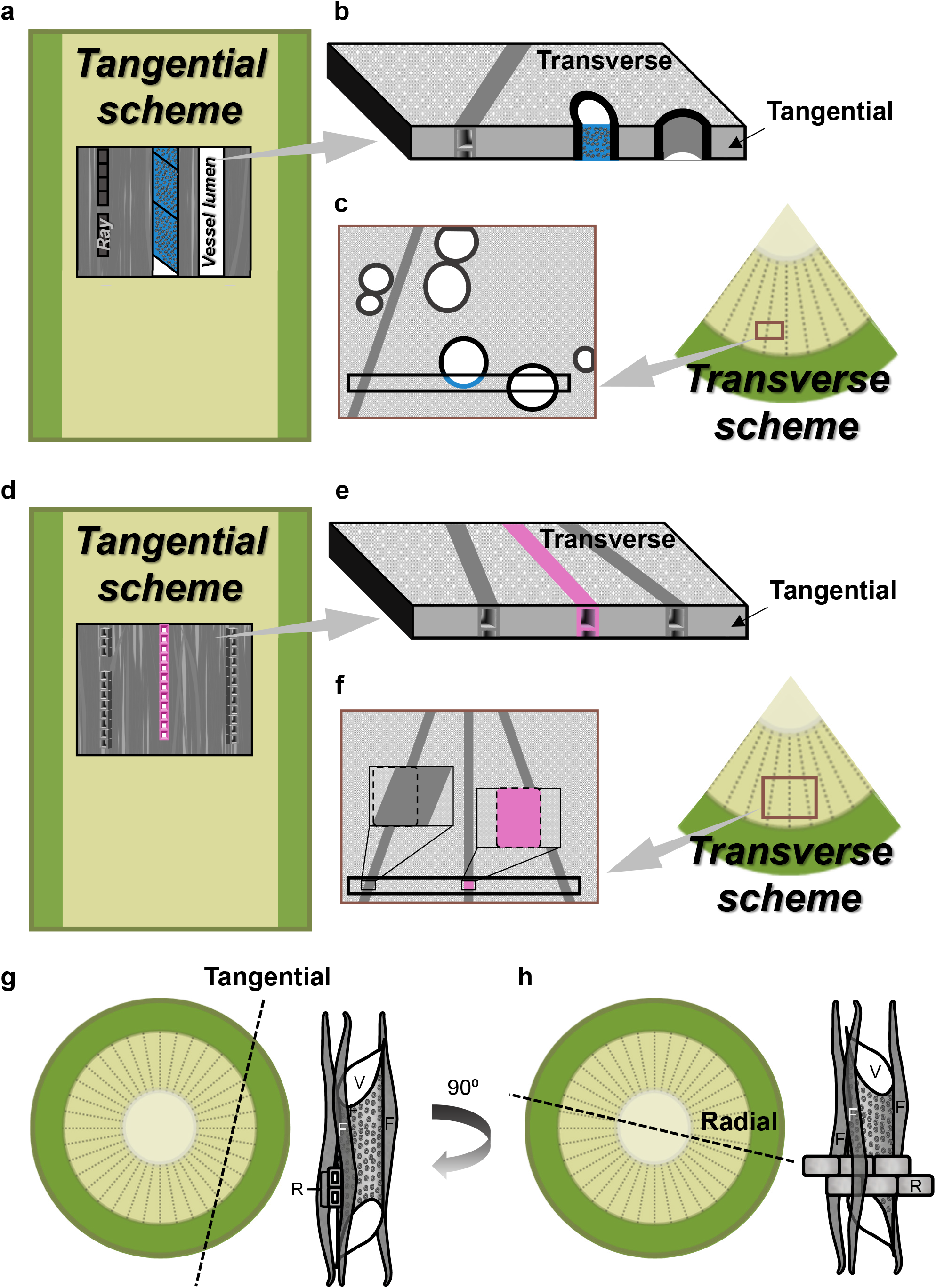
The illustrations for vessel element and ray parenchyma cell collection from transverse and tangential perspectives. **a**–**c**, The schematics of vessel elements from transverse and tangential perspectives. The blue area represents the vessel elements collected using LCM. **a**, The corresponding figure of Fig. 1i represents a tangential section with an area highlighted (blue color) for vessel element collection. **b**, A 3D structure of the highlighted area shows the location of the collected vessel elements in the stem structure schematic. **c**, The corresponding figure of Fig. 1h represents a transverse section with highlighted area. **d**–**f**, The schematics of ray parenchyma cells from transverse and tangential perspectives. The pink area represents the ray parenchyma cells collected using LCM. **d**, The corresponding figure of Fig. 1m represents a tangential section with an highlighted area (pink color). **e**, A 3D structure of the highlighted area shows the location for the collected ray parenchyma cells in the stem structure schematic. **f**, The corresponding figure of Fig. 1l represents a transverse section of the highlighted area. **g**, **h**, The 3D arrangement of three xylem cell types from tangential (**g**) and radial (**h**) perspectives. F, libriform fibers. V, vessel elements. R, ray parenchyma cells.

**Extended Data Fig. 5.**
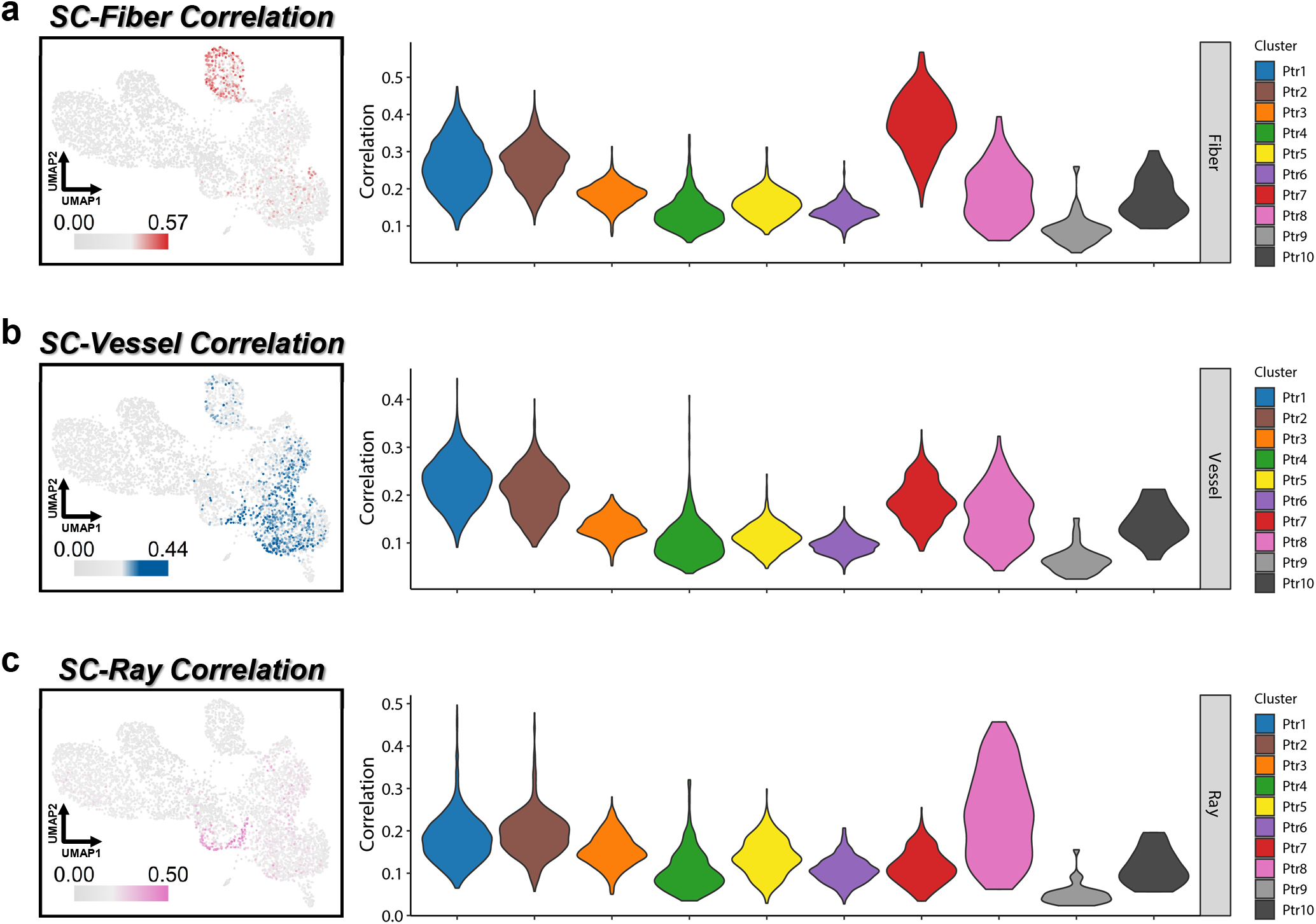
Violin plots of the transcriptomic correlation between each cell type of lcmRNA-seq and each cell cluster of scRNA-seq results. **a**–**c**, The correlations between scRNA-seq results with libriform fibers (**a**), vessel elements (**b**) or ray parenchyma cells (**c**), respectively.

**Extended Data Fig. 6.**
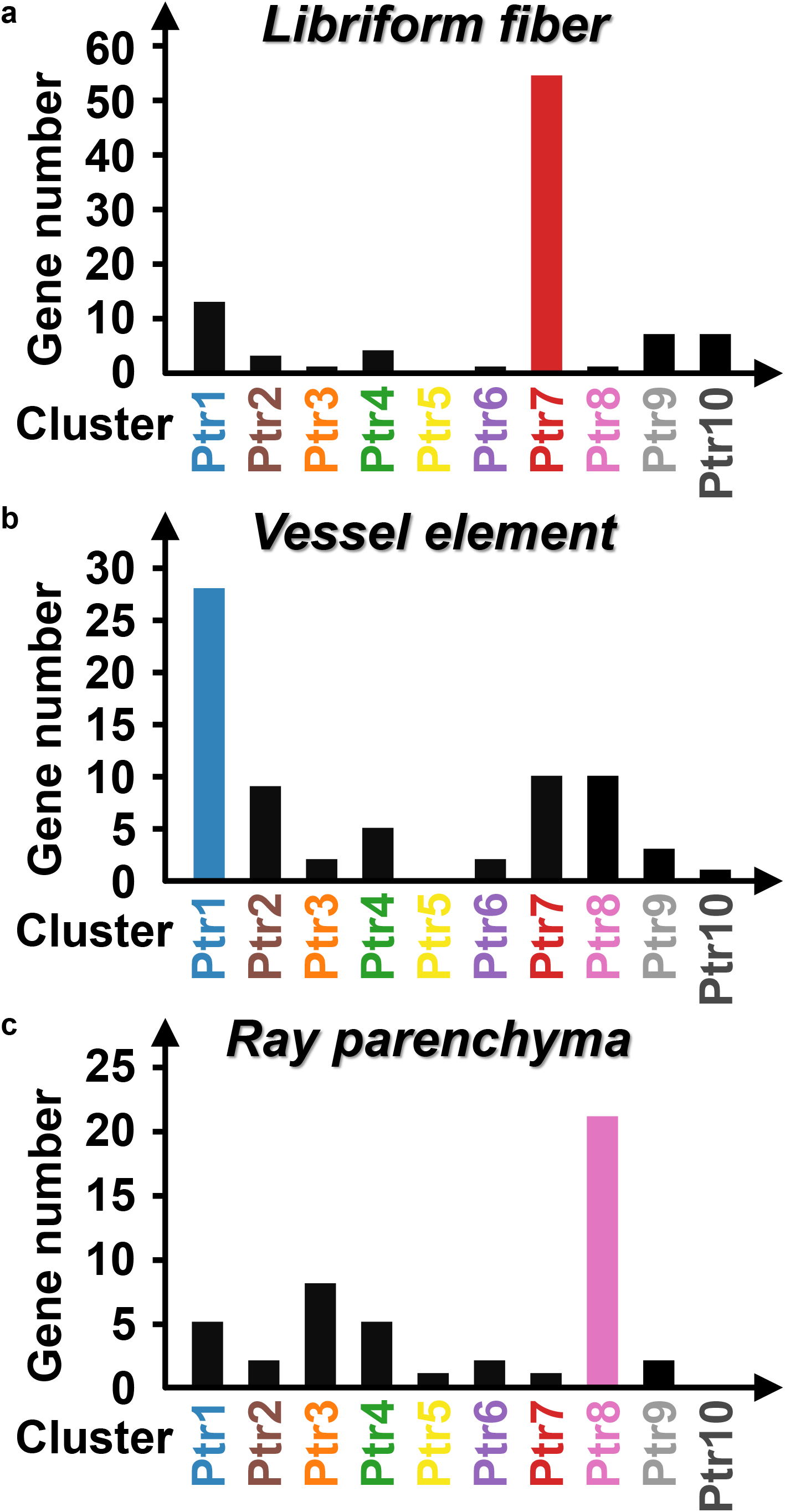
Histograms of the scUPlcmUP gene numbers. **a**–**c**, The scUPlcmUP gene numbers of libriform fibers (**a**), vessel elements (**b**) and ray parenchyma cells (**c**). Only the lcmUP genes with average transcripts per million (TPM) more than 4 were included.

**Extended Data Fig. 7.**
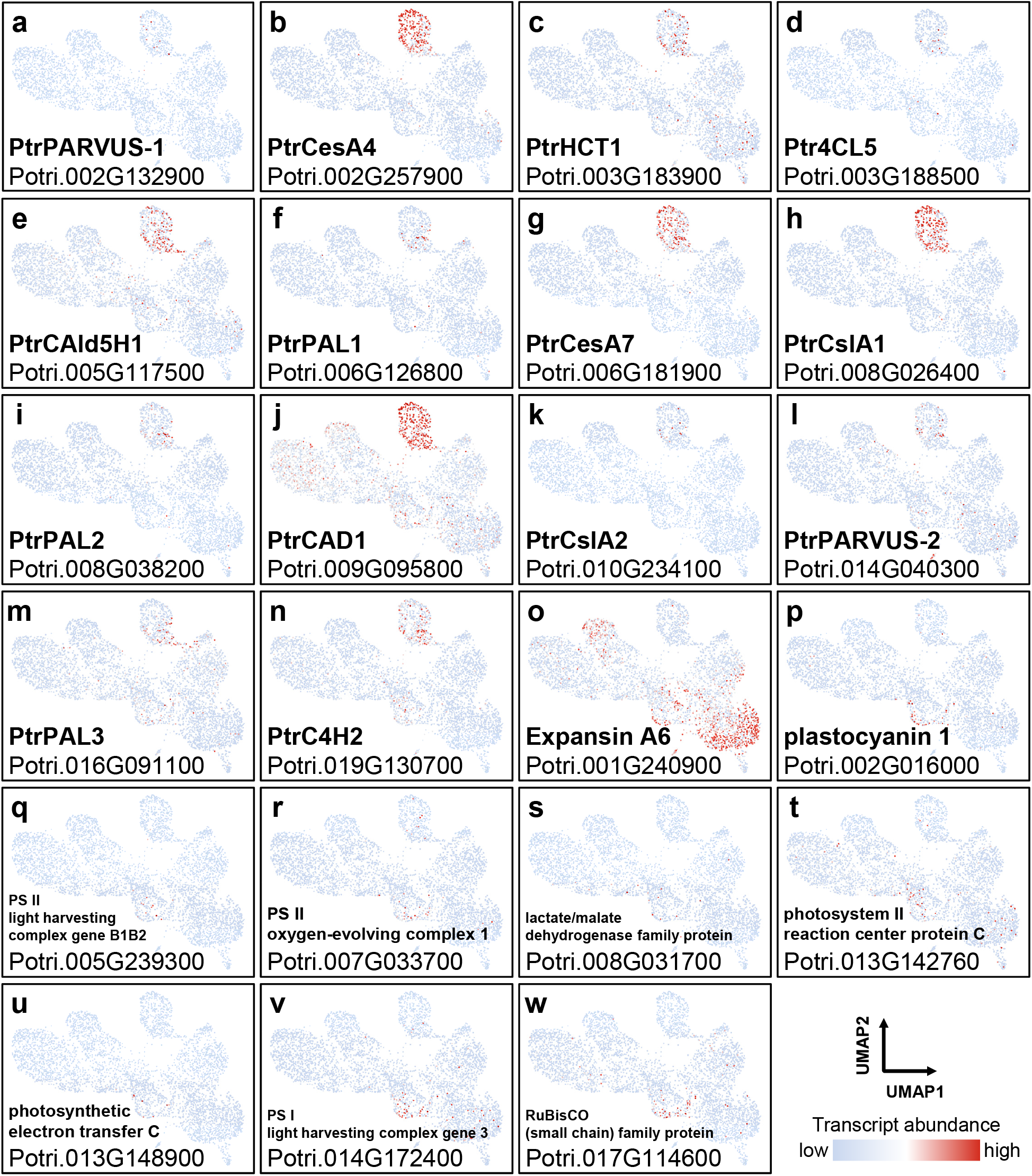
Previous known genes in the scUPlcmUP genes. **a**–**n**, Secondary biosynthesis genes in libriform fibers. **o**, The expansin gene in vessel elements. **p**–**w**, The photosynthesis-related genes in ray parenchyma cells.

**Extended Data Fig. 8.**
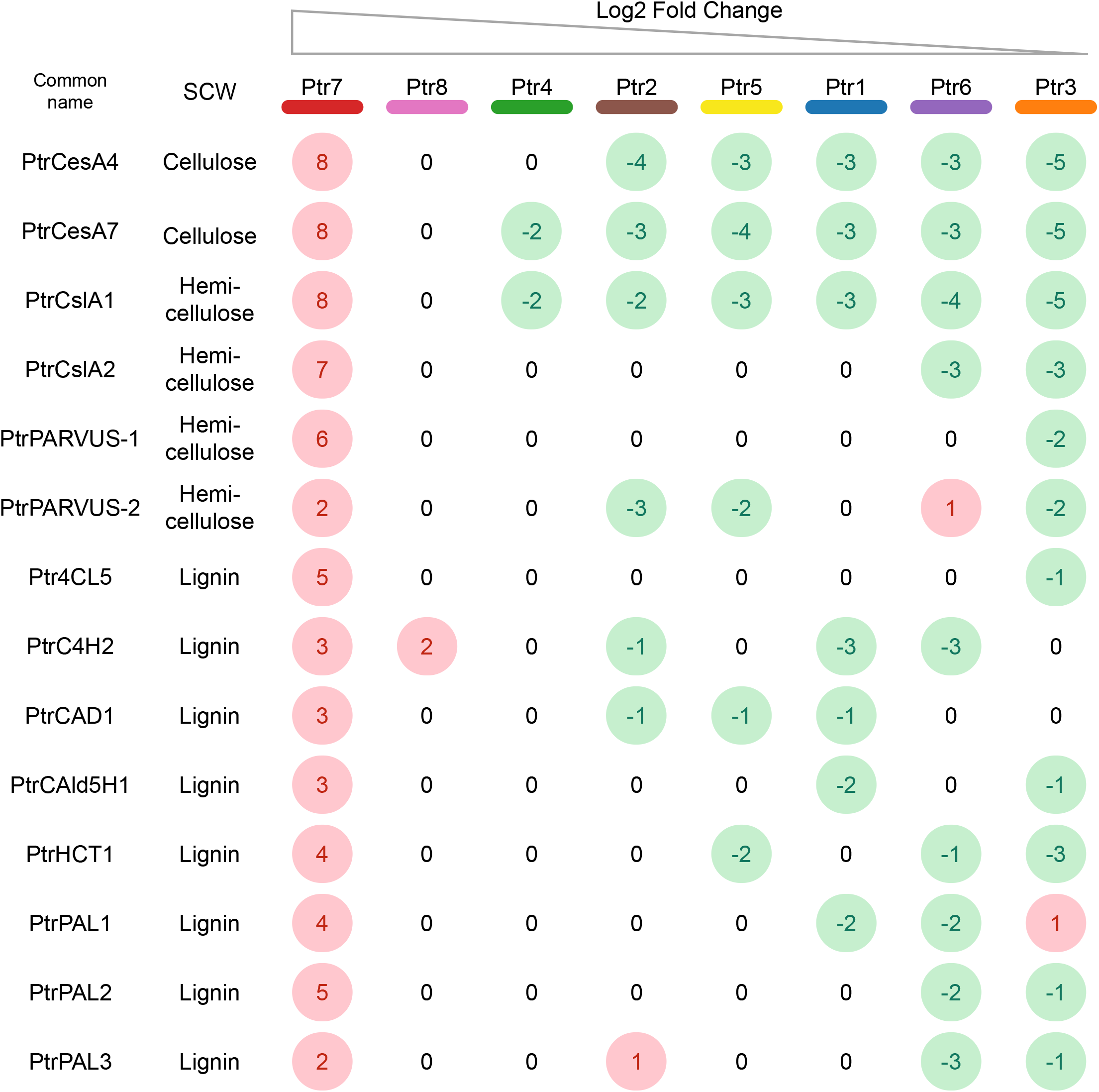
Relative transcript abundance of the secondary cell wall biosynthesis genes in the cell clusters. The relatively up- or down-regulated genes are marked as red and green colors, respectively.

**Extended Data Fig. 9.**
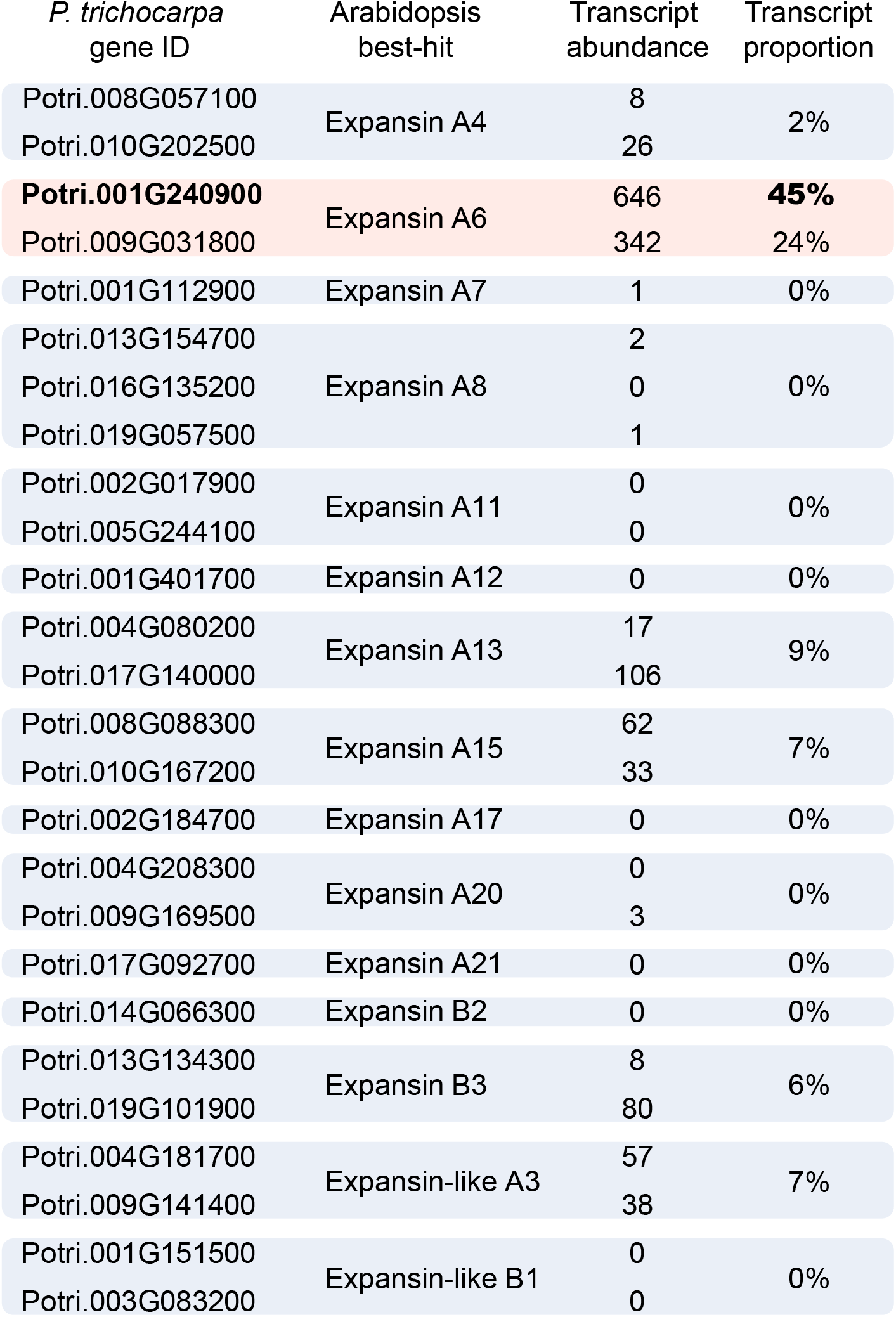
Transcript abundance of the *P. trichocarpa* homologous expansin genes of *A. thaliana*. The *P. trichocarpa* homologous expansin genes of *A. thaliana* are listed with their corresponding transcript abundance and their transcript proportion in SDX among all expansin transcript abundance.

**Extended Data Fig. 10.**
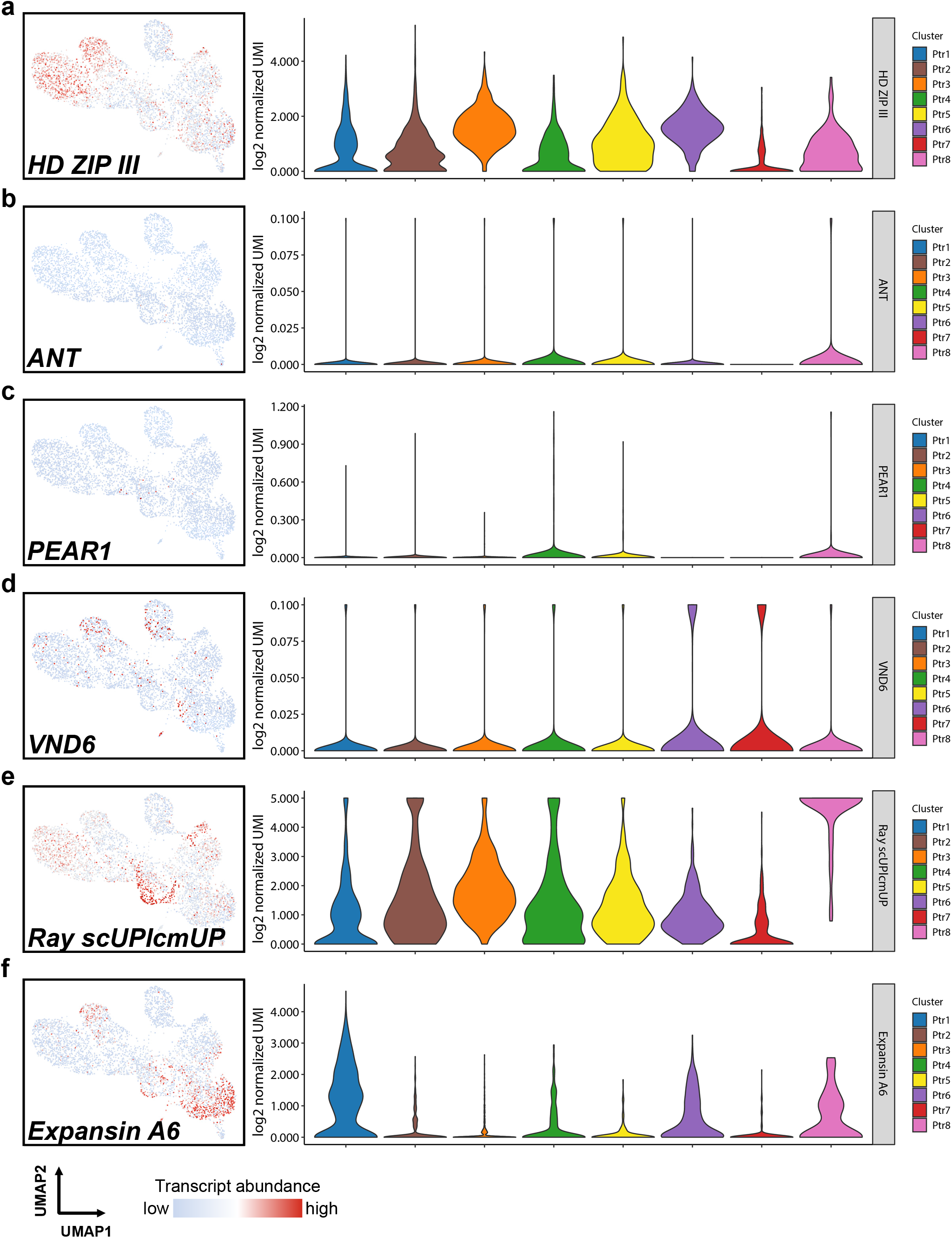
Violin plots of the transcript abundance of each orthologous group in the scRNA-seq results. **a**–**f**, Violin plots are used to reveal the log_2_ normalized UMI counts of each orthologous group in each cell cluster.

**Extended Data Fig. 11.**
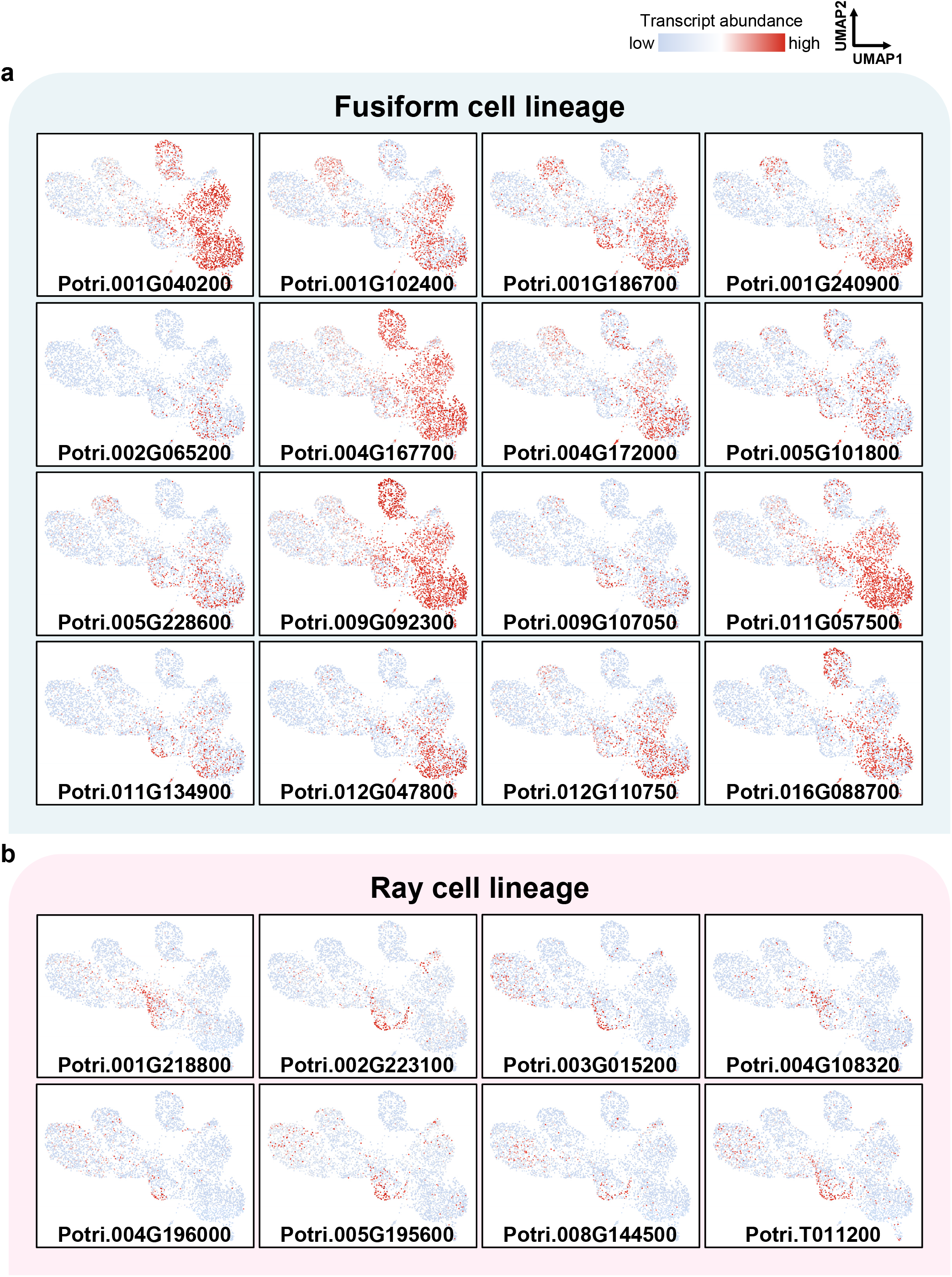
Transcript abundance of up-regulated genes in scRNA-seq results of vessel elements and ray parenchyma cells reveals fusiform and ray cell lineages. **a**, **b**, The gene expression profiles of the up-regulated genes in vessel elements and ray parenchyma cells on the UMAP show continuous cell lineages for fusiform (**a**) and ray cells (**b**), respectively.

**Extended Data Fig. 12.**
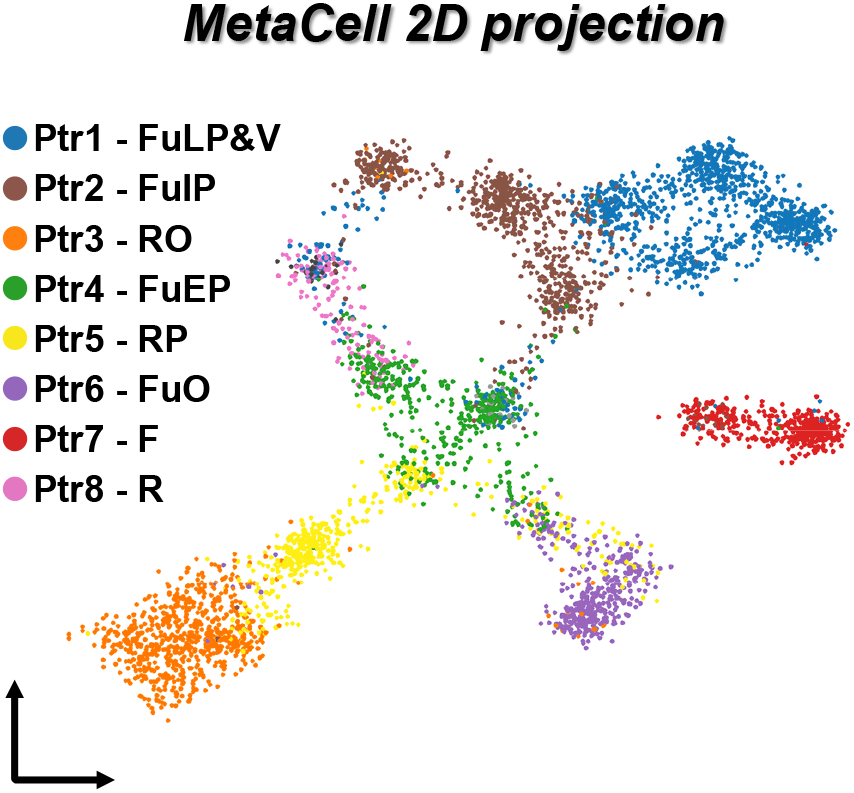
MetaCell plot of the cell clusters in *P. trichocarpa*. 2D projection of *P*. *trichocarpa* SDX single cells using MetaCell dimensionality reduction. Cells are labeled with colors corresponding to those in Fig. 1b. FuLP&V, fusiform late precursor and vessel element. FuIP, fusiform intermediate precursor. RO, ray organizer. FuEP, fusiform early precursor. RP, ray precursor. FuO, fusiform organizer. F, libriform fiber. R, ray parenchyma cell.

**Extended Data Fig. 13.**
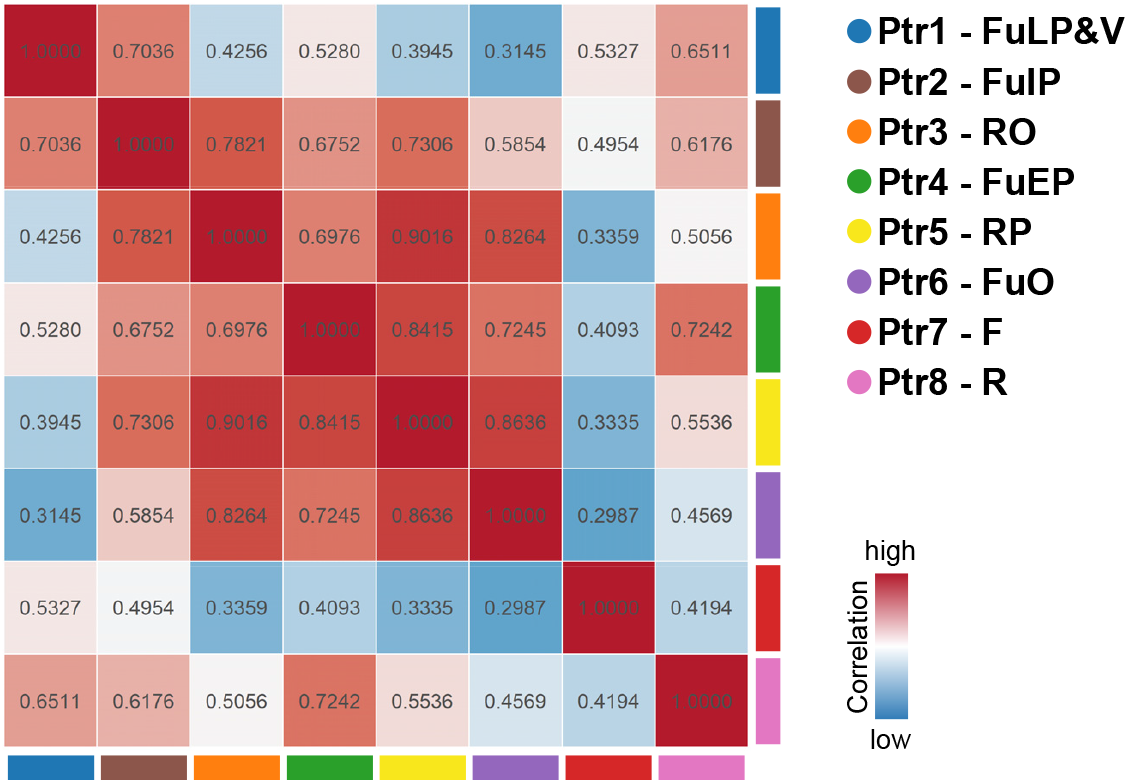
Pairwise correlation of the cell clusters in *P. trichocarpa*. Pairwise correlation between transcriptomic profiles of Ptr1–Ptr8 of *P. trichocarpa* SDX. FuLP&V, fusiform late precursor and vessel element. FuIP, fusiform intermediate precursor. RO, ray organizer. FuEP, fusiform early precursor. RP, ray precursor. FuO, fusiform organizer. F, libriform fiber. R, ray parenchyma cell.

**Extended Data Fig. 14.**
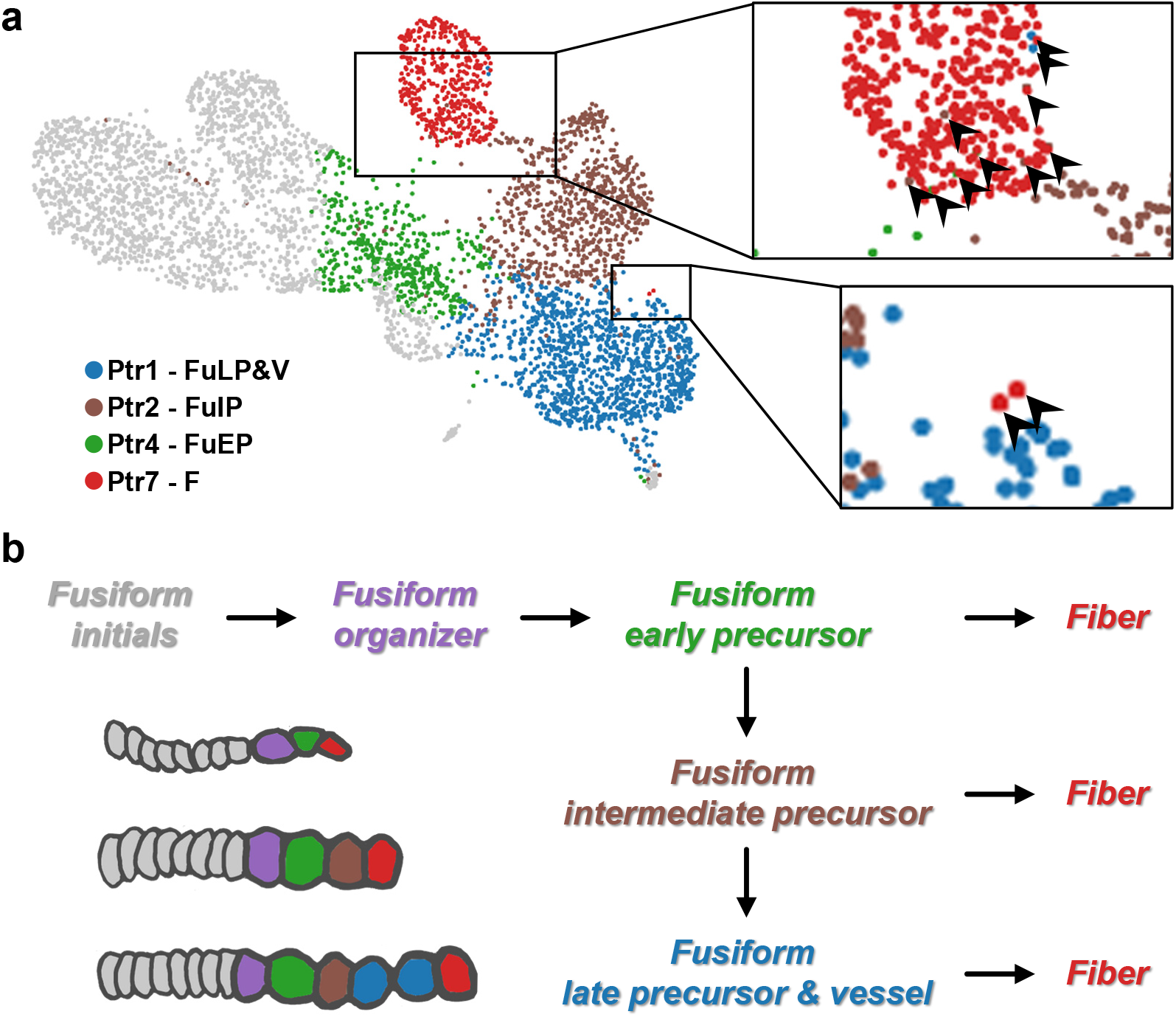
Fusiform cell clusters reveal the potency to transit into libriform fibers in *P. trichocarpa*. **a**, The fusiform cell clusters on UMAP plot and the area within the black boxes are magnified. The black arrows indicate the intermixing cells between libriform fibers (F, Ptr7), early fusiform precursors (FuEP, Ptr4), intermediate fusiform precursors (FuIP, Ptr2) and late fusiform precursors/vessel elements (FuLP&V, Ptr1). **b**, Schematics of libriform fiber cell lineages in SDX.

**Extended Data Fig. 15.**
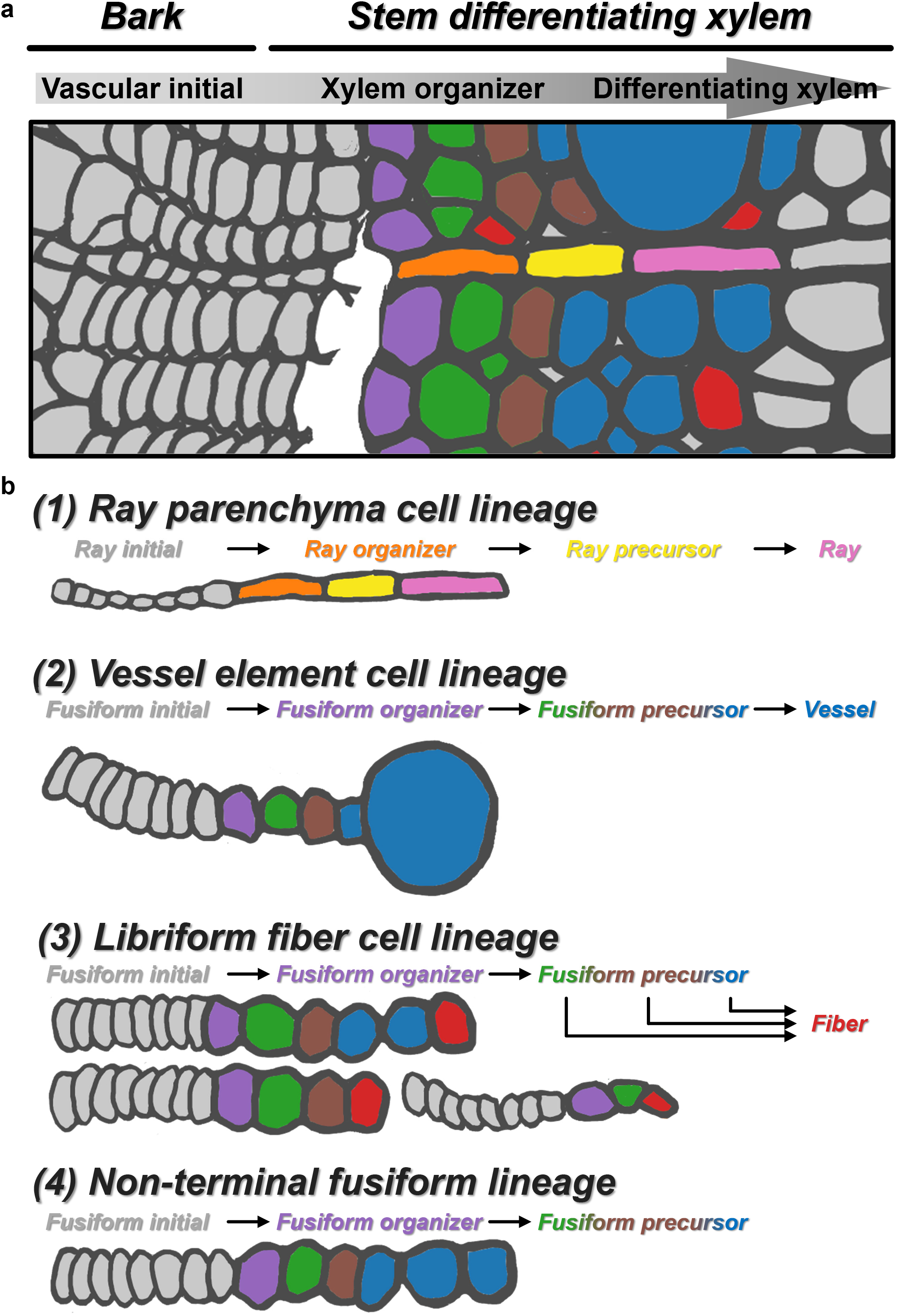
Schematics of different developmental cell lineages in SDX in *P. trichocarpa*. **a**, After stem debarking, the revealed SDX on the stem surface contains xylem organizer and differentiating xylem, which includes different cell lineages. **b**, Three types of cell lineages in SDX are ray parenchyma cell, vessel element and libriform fiber cell lineages. Another incomplete or undergoing cell lineage is shown as non-terminal fusiform lineage. All lineages start with initials to precursors then to terminal differentiated ray parenchyma cells, vessel elements or libriform fibers.

**Extended Data Fig. 16.**
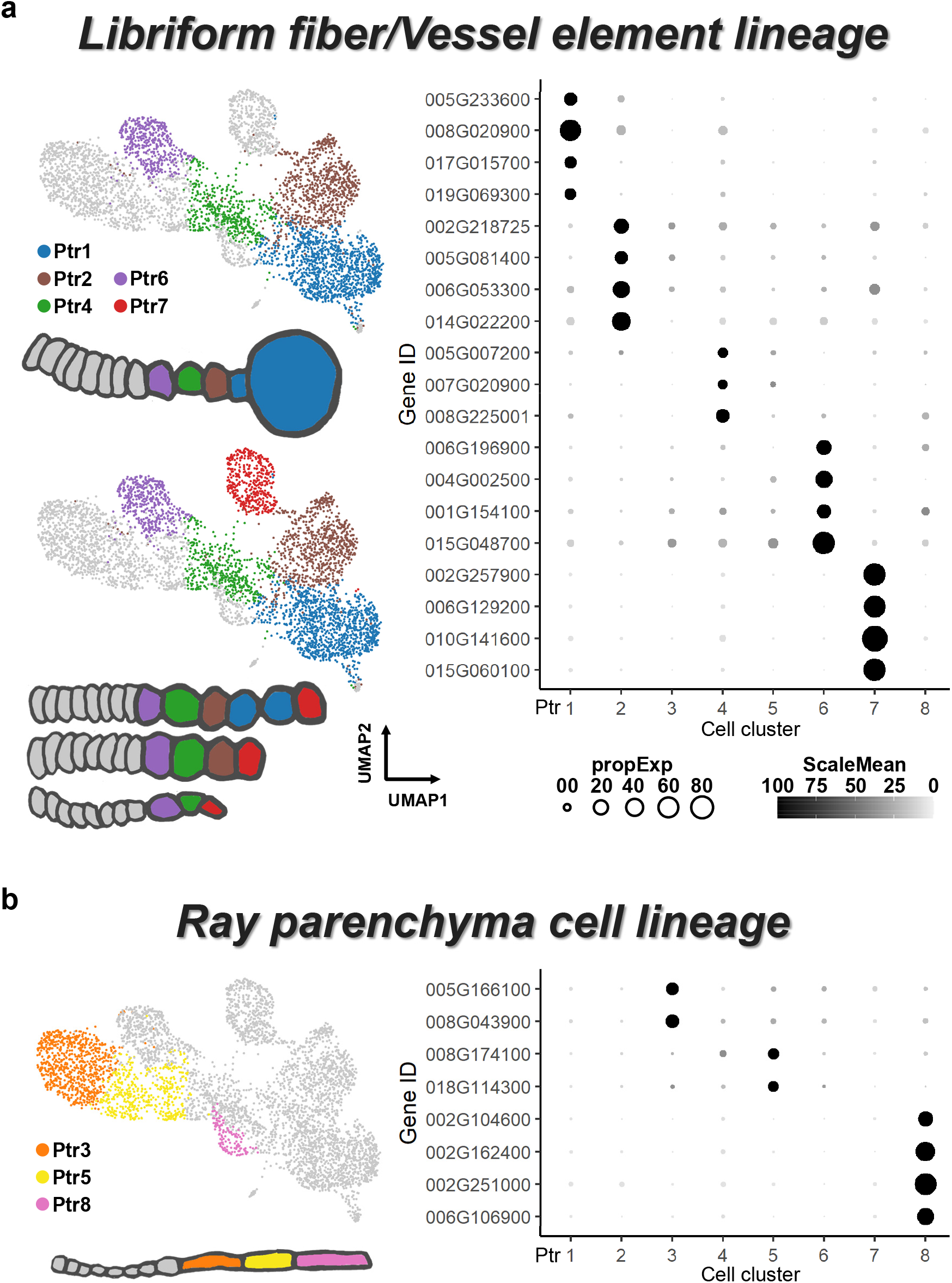
Dot plots exhibit preferential patterns of marker genes in each cell cluster of libriform fiber/vessel element or ray parenchyma cell lineages. **a, b,** The dot size represents the proportion of cells in each cell cluster with the marker gene expression, and the dot brightness shows the relative mean gene expression of the marker genes in libriform fiber/vessel element (**a**) or ray parenchyma cell lineages (**b**), respectively.

**Extended Data Fig. 17.**
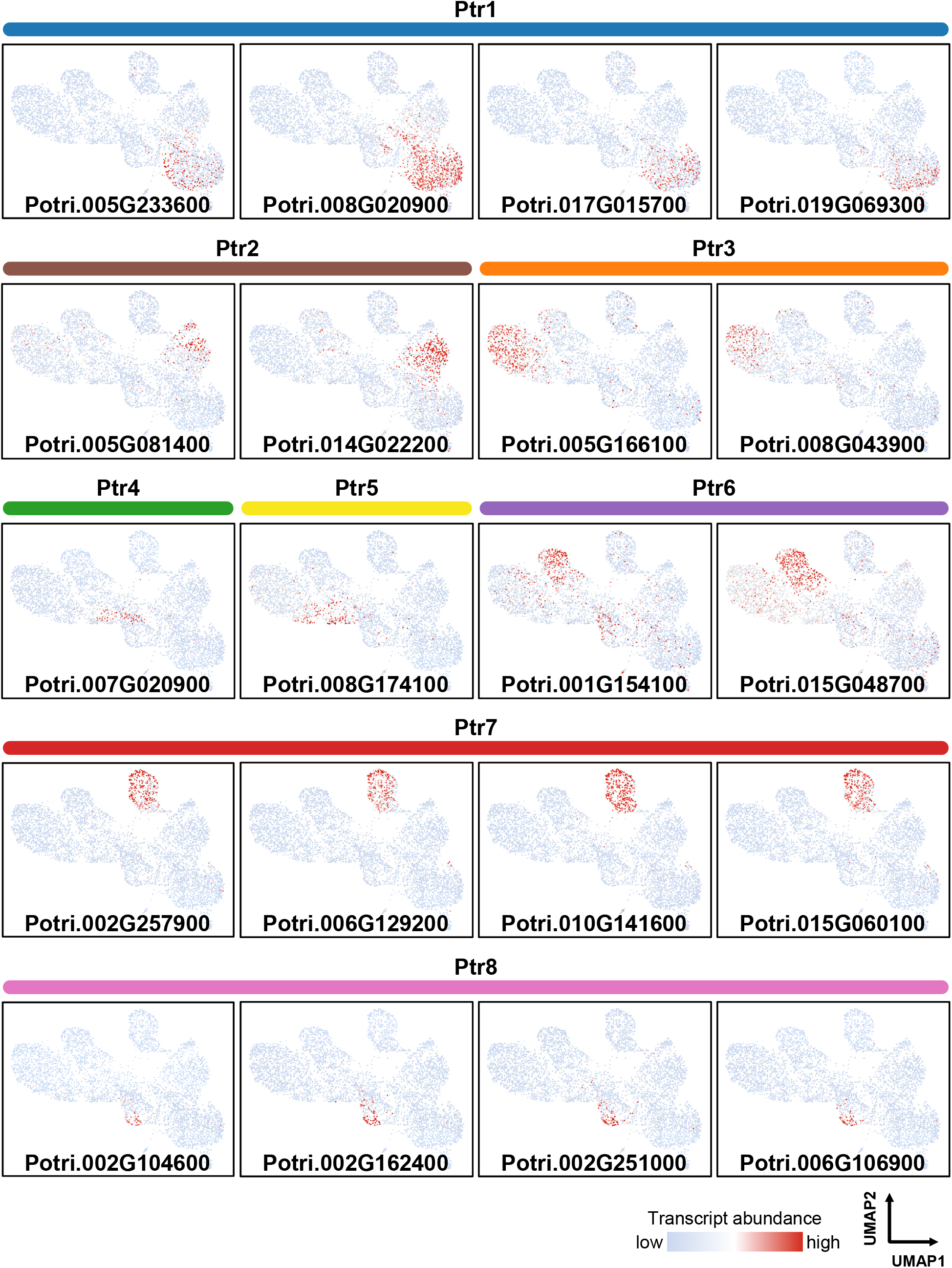
Transcript abundance of marker genes of each cell cluster in *P. trichocarpa*. Many DEGs from each cell cluster are identified as marker genes if these genes show an exclusive expression in that cell cluster.

**Extended Data Fig. 18.**
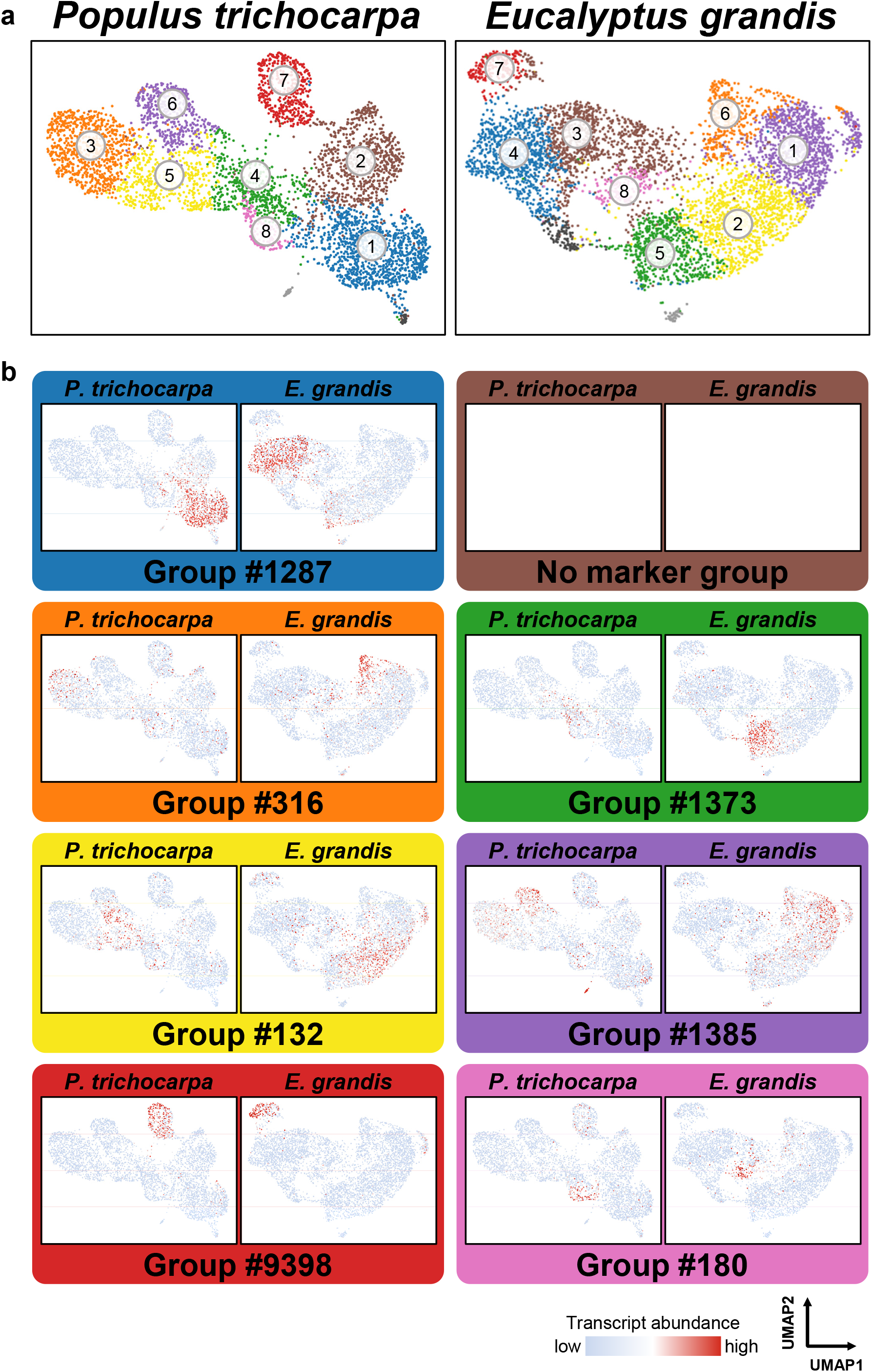
Representative marker orthologous groups exclusively expressed in the different cell clusters in both *P. trichocarpa* and *E. grandis*. **a**, The cell cluster plots obtained through unsupervised K-means clustering and UMAP are based on the SDX scRNA-seq results of *P. trichocarpa* and *E. grandis*, respectively. **b**, The cluster-exclusive distributions of each marker orthologous group are represented by Group #1287 for vessel element/late fusiform precursor (blue), Group #316 for ray organizer (orange), Group #1373 for fusiform early precursor (green), Group #132 for ray precursor (yellow), Group #1385 for fusiform organizer (purple), Group #9398 for libriform fiber (red) and Group #180 for ray parenchyma cell (pink). No marker orthologous groups are observed in fusiform intermediate precursor (brown).

**Extended Data Fig. 19.**
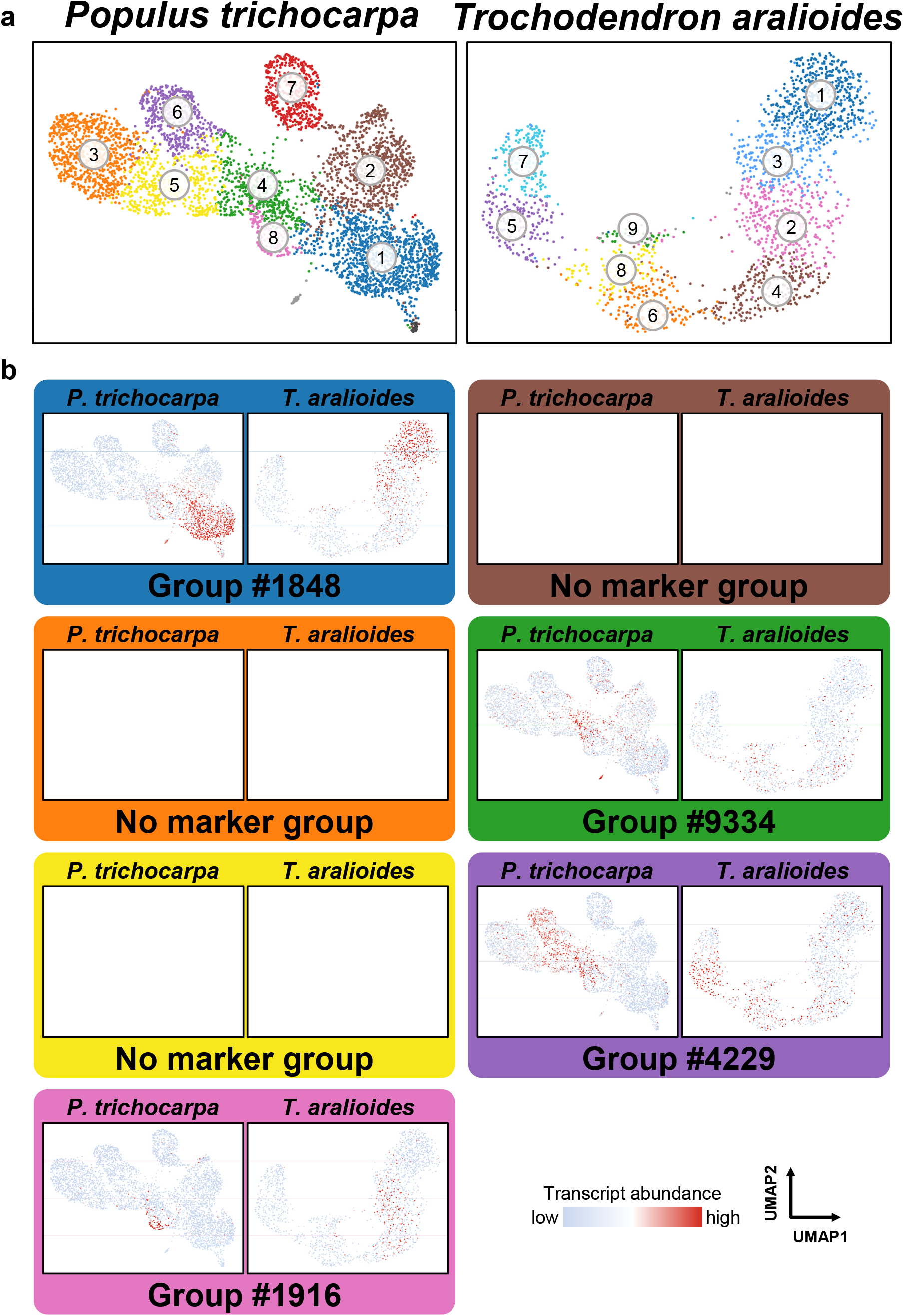
Representative marker orthologous groups exclusively expressed in the different cell clusters both in *P. trichocarpa* and *T. aralioides*. **a**, The cell cluster plots obtained through unsupervised K-means clustering and UMAP are based on the SDX scRNA-seq results of *P. trichocarpa* and *T. aralioides*, respectively. **b**, The cluster-exclusive distributions of each marker orthologous group are represented by Group #1848 for vessel element/late fusiform precursor (blue), Group #9334 for fusiform early precursor (green), Group #4229 for fusiform organizer (purple) and Group #1916 for ray parenchyma cell (pink). No marker orthologous groups are observed in fusiform intermediate precursor (brown), ray organizer (orange) and ray precursor (yellow).

**Extended Data Fig. 20.**
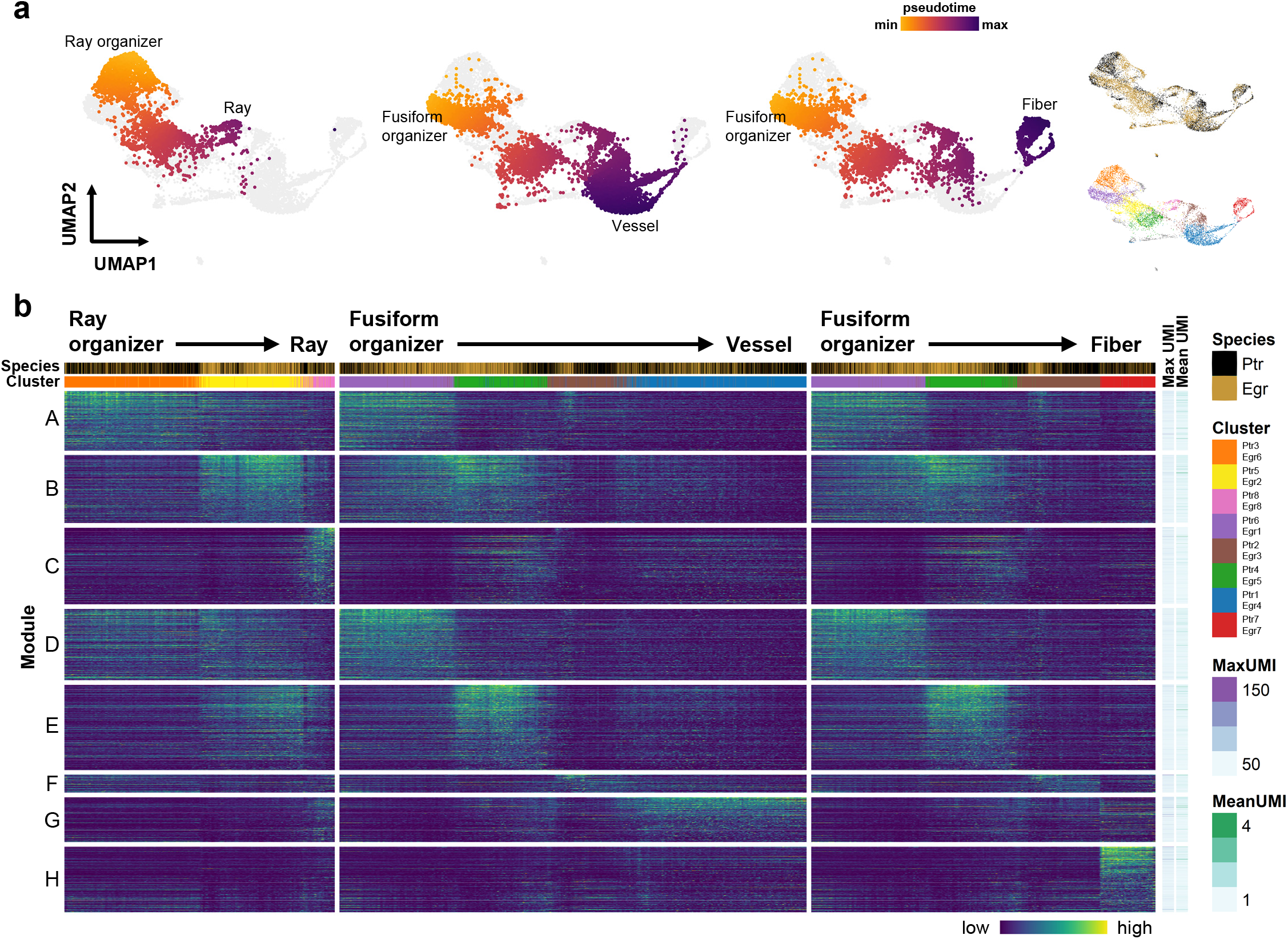
Pseudotime analysis to reveal the temporal expression pattern of each cell lineage using two-species clustering of *P. trichocarpa* and *E. grandis*. **a**, Ray parenchyma, vessel element and libriform fiber lineages are from ray organizer or fusiform organizer. **b**, Representative orthologous groups for each cell cluster can be categorized into eight modules (A–H). Along with the lineages two color bars represent as cells from two species (*P. trichocarpa* as black and *E. grandis* as gold) and their corresponding clusters. Maximum and mean UMI counts of each orthologous group are shown.

**Extended Data Fig. 21.**
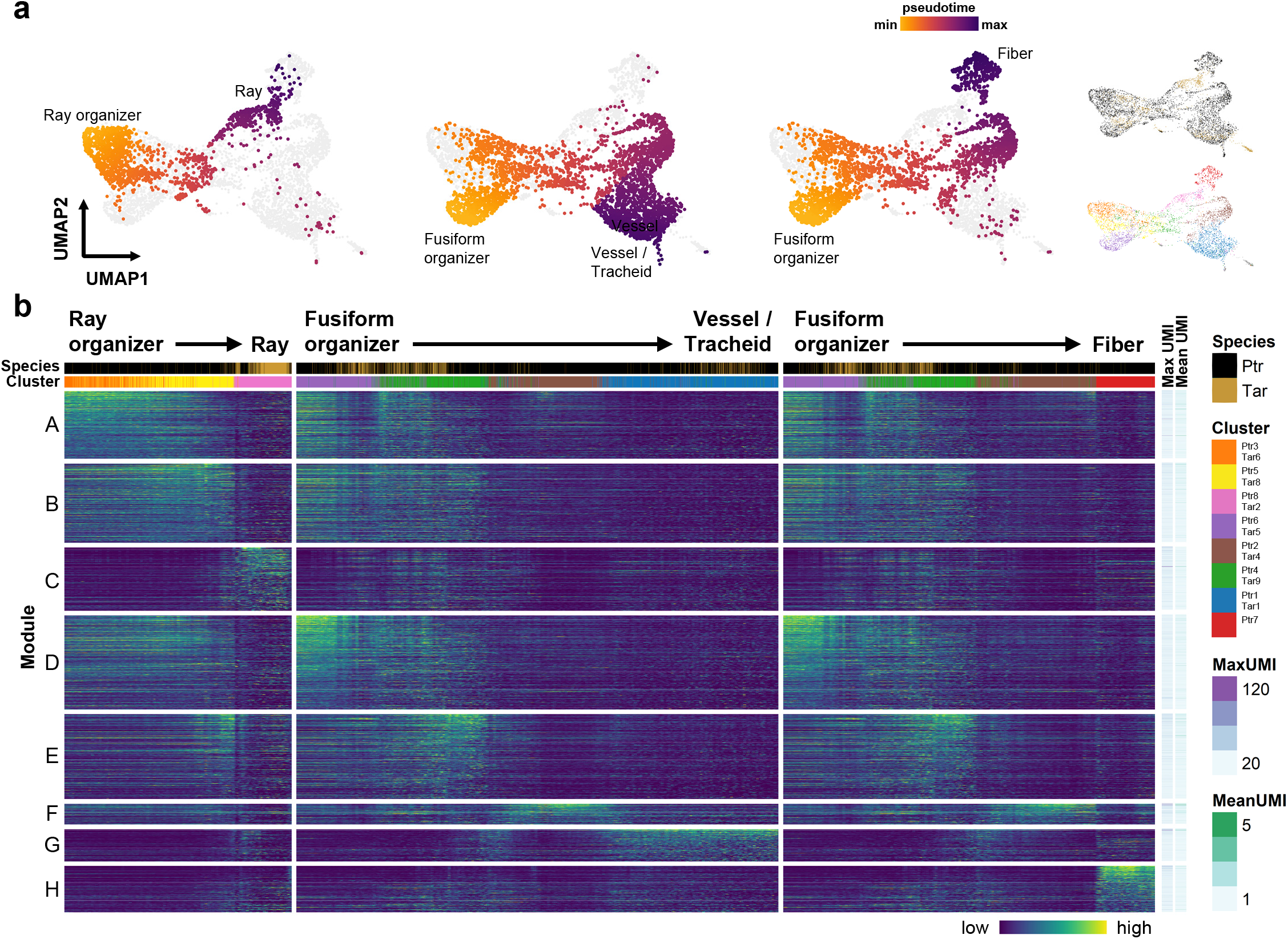
Pseudotime analysis to reveal the temporal expression pattern of each cell lineage using two-species clustering of *P. trichocarpa* and *T. aralioides*. **a**, Ray parenchyma, vessel element/tracheid and libriform fiber lineages from ray organizer or fusiform organizer are shown. **b**, Representative orthologous groups for each cell cluster can be categorized into eight modules (A–H). Along with the lineages two color bars represent as cells from two species (*P. trichocarpa* as black and *T. aralioides* as gold) and their corresponding clusters. Only *P. trichocarpa* exhibits cells in libriform fiber cluster (Ptr7) of the libriform fiber lineage. Maximum and mean UMI counts of each orthologous group are shown.

**Extended Data Fig. 22.**
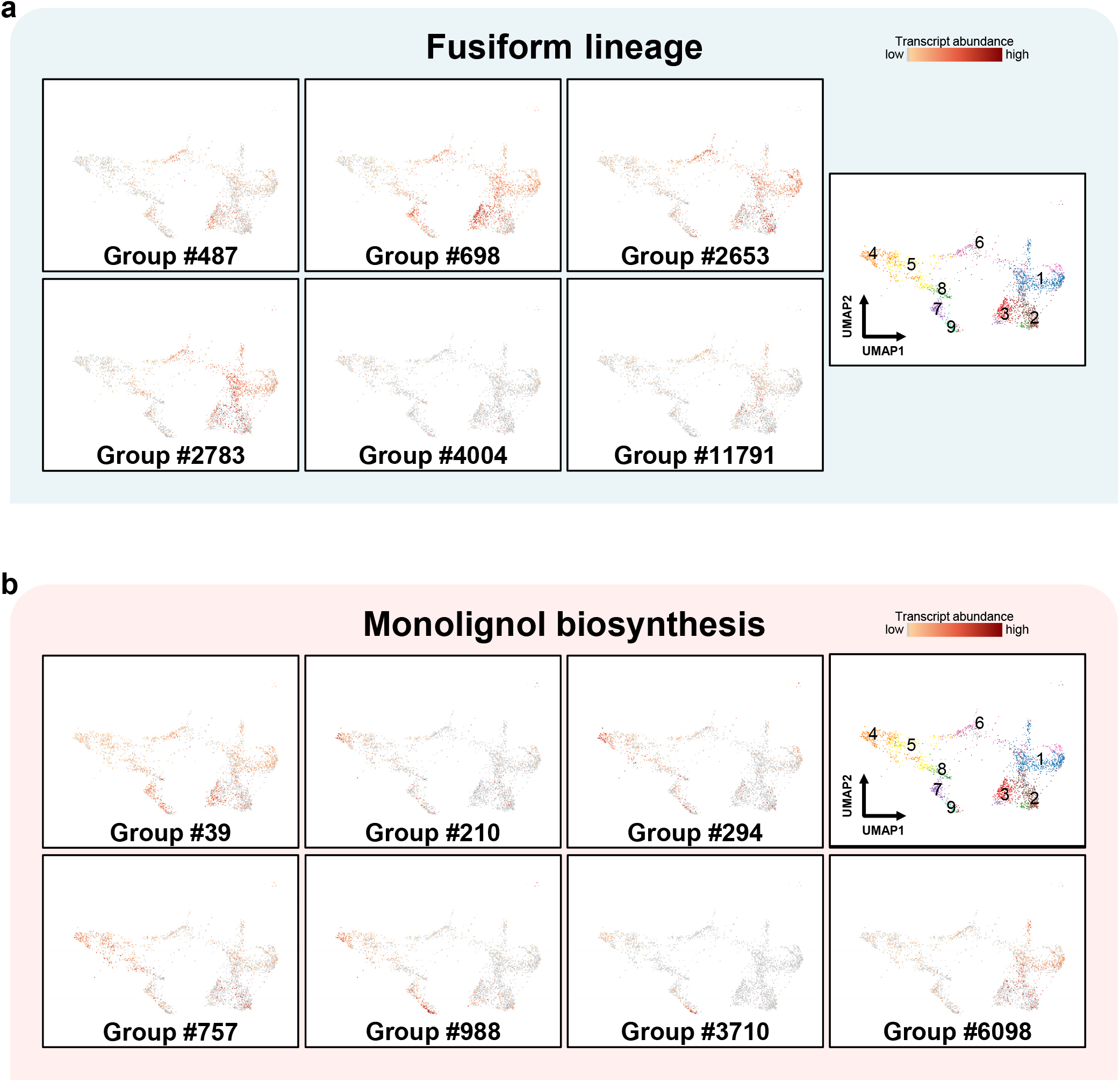
The *L. chinense* homologous genes of the fusiform marker genes and monolignol biosynthesis genes from *P. trichocarpa*. **a**, **b**, The transcript abundance of the *L. chinense* homologous genes of the fusiform marker genes (**a**) and monolignol biosynthesis genes (**b**) from *P. trichocarpa* are shown on the UMAP plots of two-species analyses (Fig. 3c (iv)).

**Extended Data Fig. 23.**
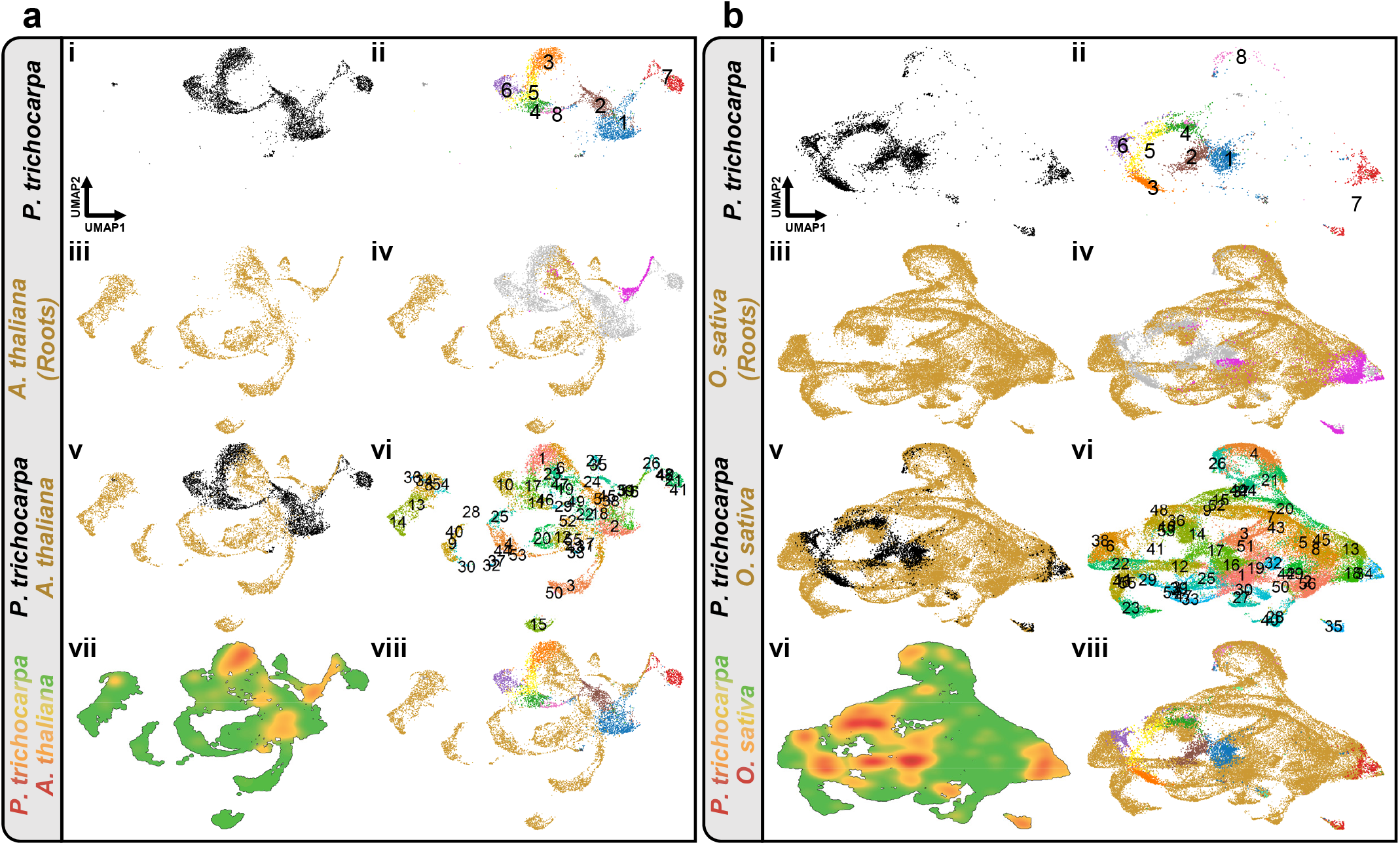
Two-species clustering and visualization of scRNA-seq data between *P. trichocarpa* and *A. thaliana* or *O. sativa*. **a**, **b**, Two-species clustering of SDX cells in *P. trichocarpa* and root cells in *A. thaliana* (**a**) or *O. sativa* (**b**). (i)–(iii), Single-species unsupervised K-means clustering. (iv)–(viii), Two-species graph-based cell clustering using orthologous genes. (i), (v), Black dots are SDX cells from *P*. *trichocarpa*. (iii), (iv), (v), (viii), Gold dots are cells from *A. thaliana* or *O. sativa*. (iv), Grey dots represent the SDX cells from *P. trichocarpa*, and the xylem cells identified in previous *Arabidopsis* or rice studies are in magenta. (ii), (viii), The colors of cell clusters are based on the single-species cell clustering results. (vi), The cell clusters in two-species clustering. (vii), The cell density of the two-species clustering. Green to orange to red colors indicate the density from low to high.

**Extended Data Fig. 24.**
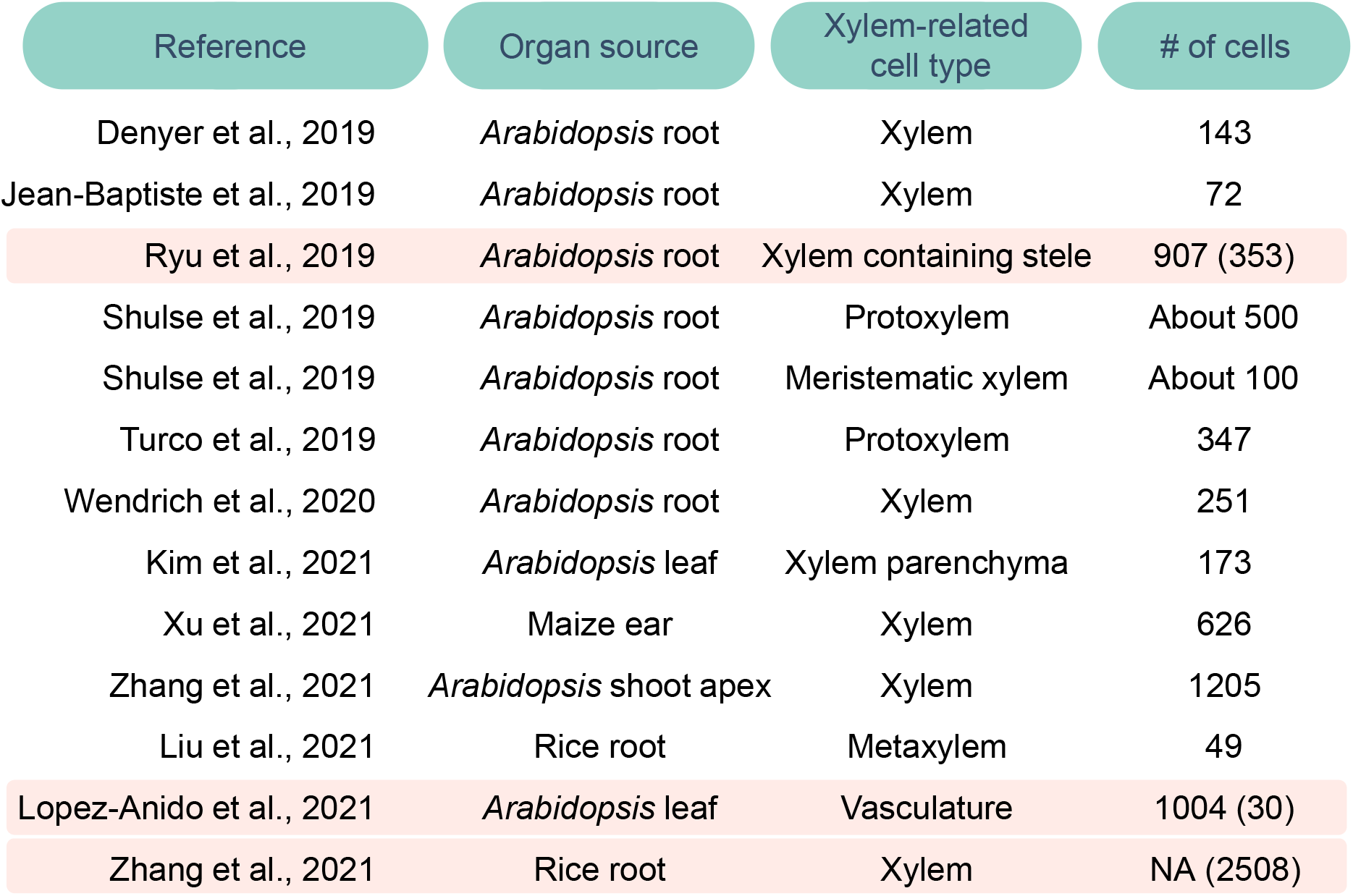
A summary of the xylem cells identified from previous studies. The previous studies with sizable cell numbers from xylem-related cell types are used in this study with highlighted light red color as the background. Three sets of cells from Ryu et al. (907 cells from xylem containing stele)^13^, Lopez-Anido et al. (1004 cells from vasculature)^22^ and Zhang et al. (xylem cell number was not available (NA))^21^ are selected. Through previously reported marker genes, total 353, 30 and 2508 xylem cells are identified.

**Extended Data Fig. 25.**
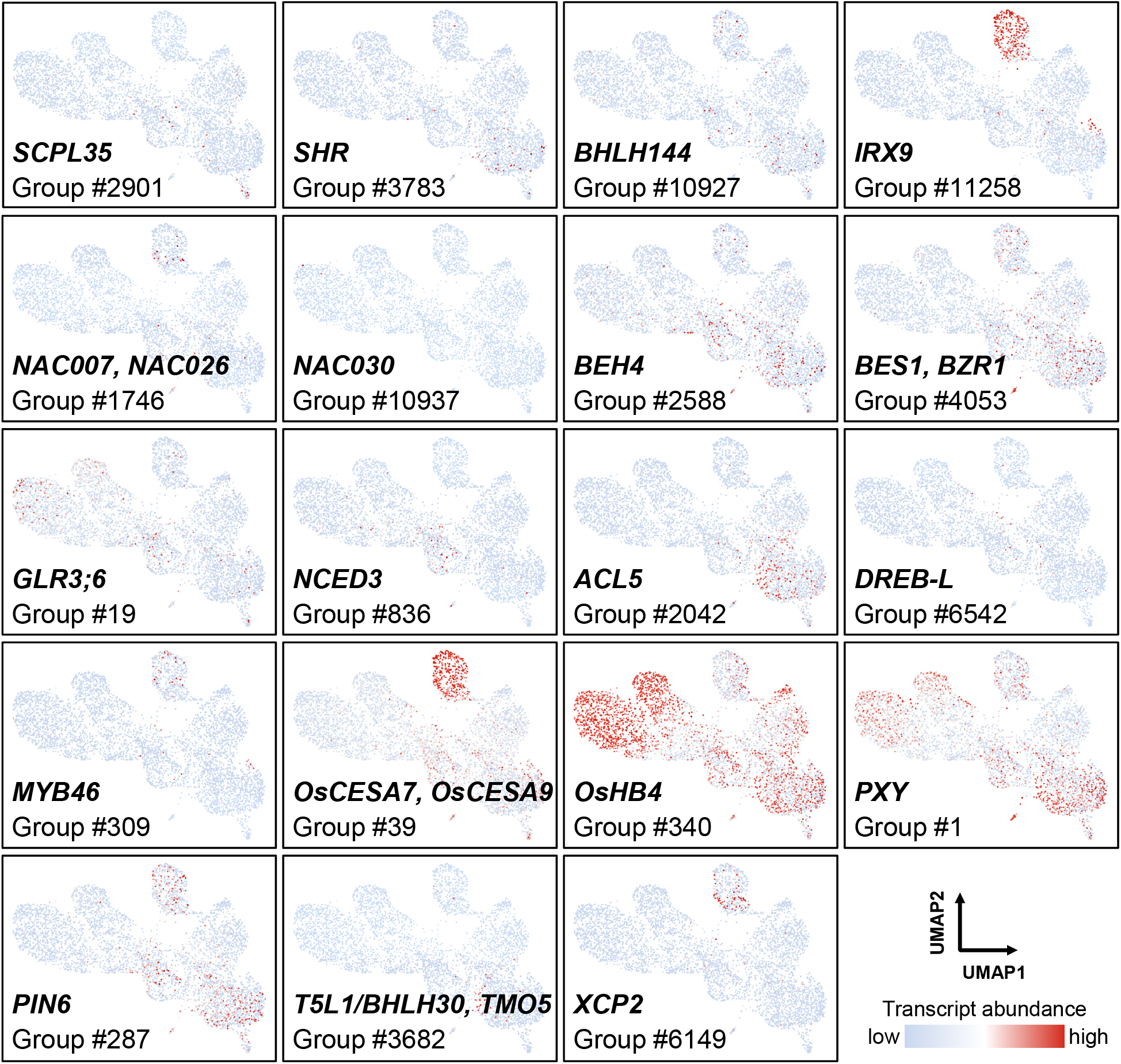
Transcript abundance of previously identified xylem marker genes. The transcript abundance of the *P. trichocarpa* homologous genes of *A. thaliana* and *O. sativa* are shown on the UMAP plots of scRNA-seq results.

**Extended Data Fig. 26.**
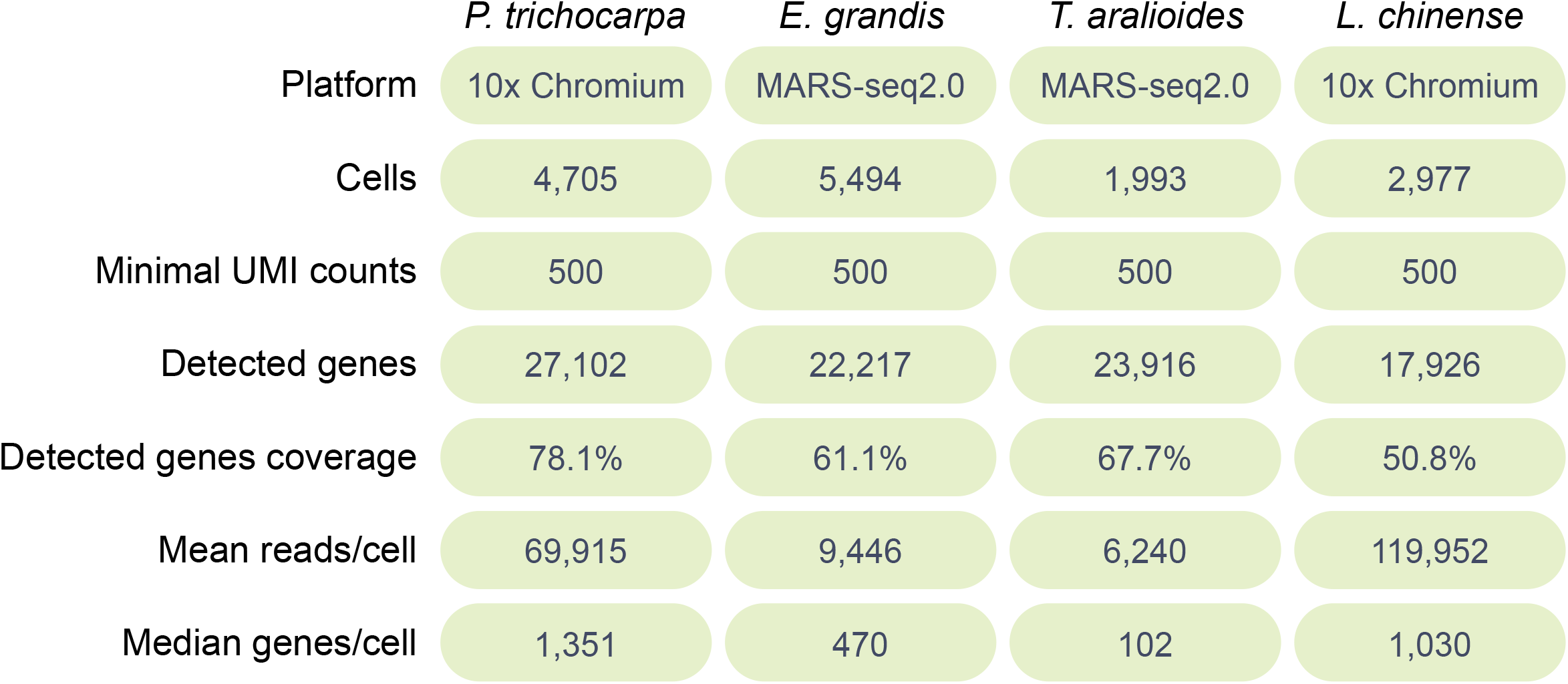
Summary of scRNA-seq assays of SDX in *P. trichocarpa*, *E. grandis*, *T. aralioides* and *L. chinense*. Statistics of scRNA-seq profiling by 10x Chromium or MARS-seq2.0 are shown for cells with at least 500 UMIs, including cell numbers, total detected genes, percentage of annotated genes that are detected, mean read counts per cell and median number of genes detected per cell.

**Extended Data Fig. 27.**
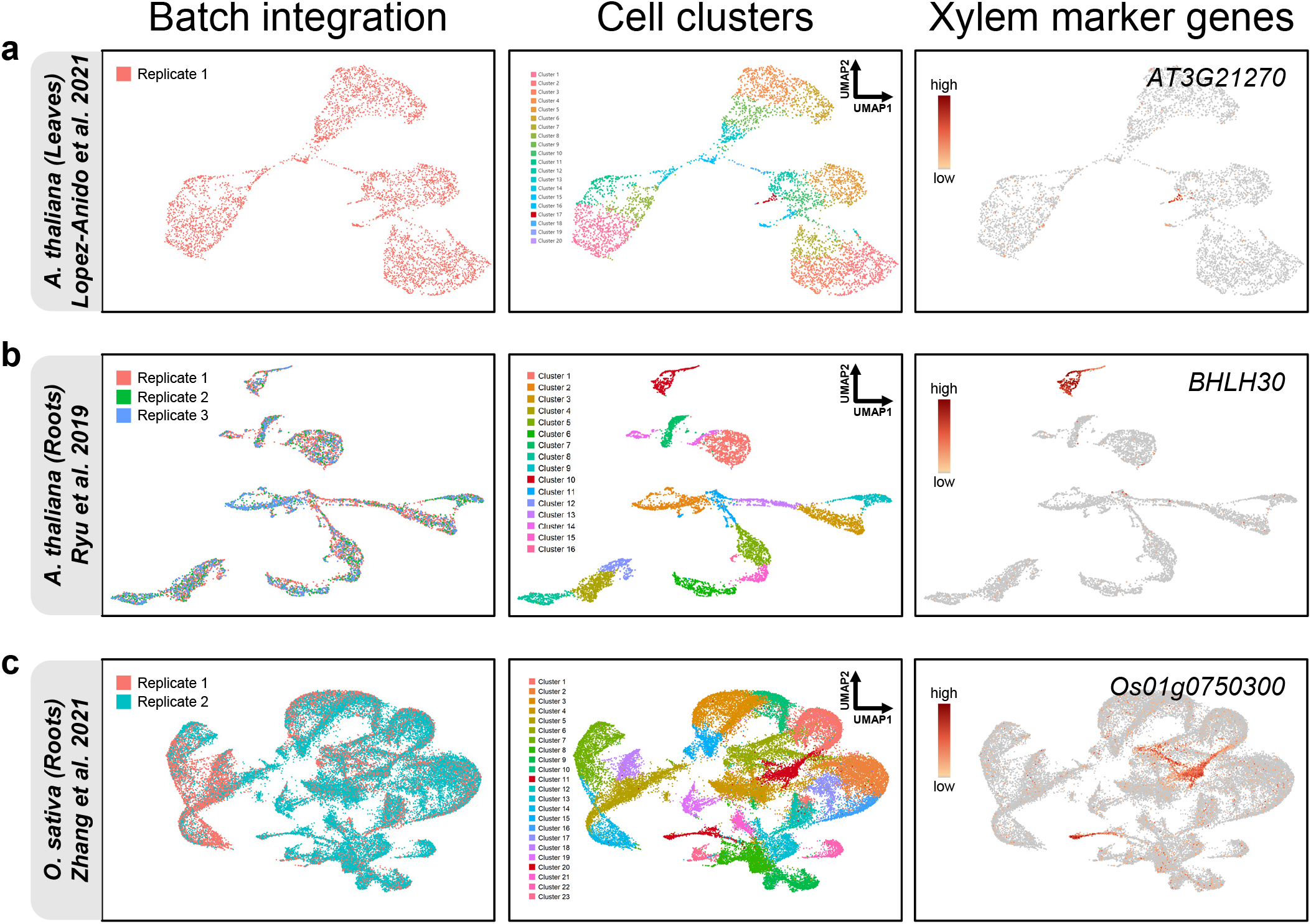
Xylem cell identification from previous scRNA-seq results. **a**–**c**, Different replicates of the scRNA-seq results from *A. thaliana* roots and *O. sativa* roots are first integrated for cell clustering using Seurat pipeline. Previous identified xylem cells using marker genes reveal that the xylem cells locate at Cluster 17, Cluster 10 and Cluster 11/20 in *A. thaliana* leaves (**a**), roots (**b**) and *O. sativa* roots (**c**), respectively.

**Extended Data Fig. 28.**
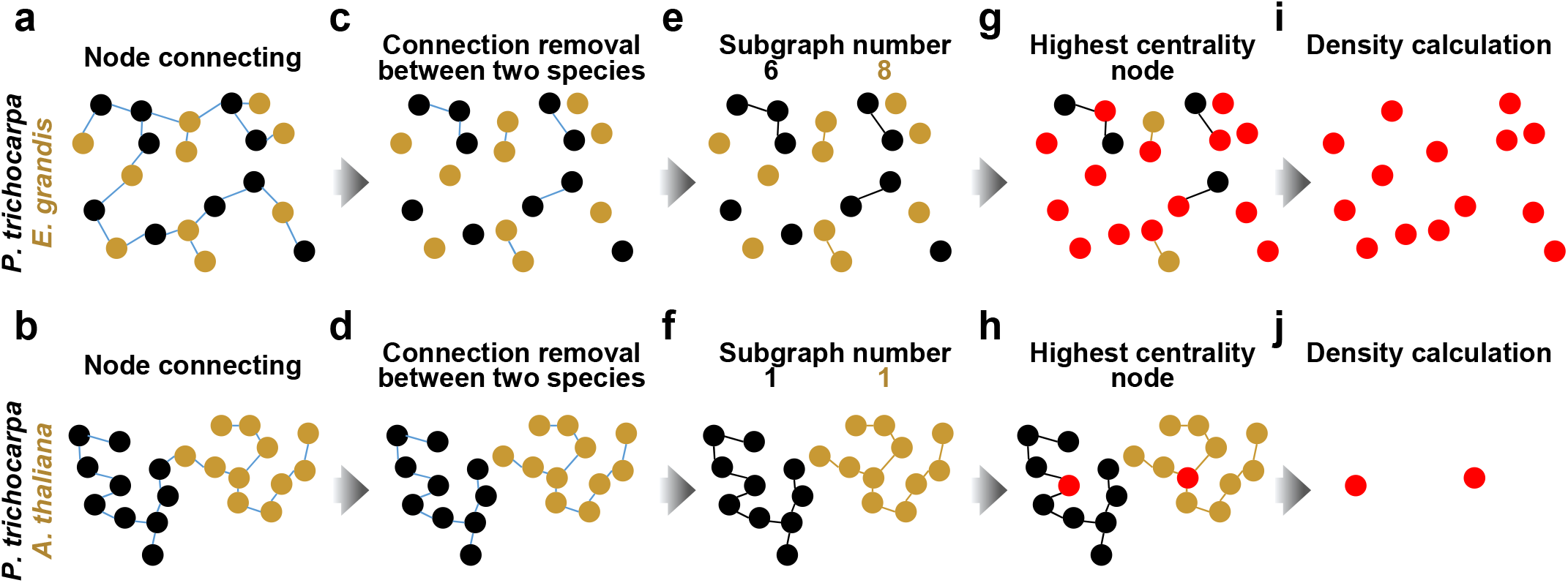
Cell density analysis of two-species clustering. Take two schematics of two-species clustering as examples, one as *P. trichocarpa* and *E. grandis* and the other one as *P. trichocarpa* and *A. thaliana*. **a**, **b**, All the nodes as cells from two species are connected with the shortest line length. **c**, **d**, The lines between the cells from different species are removed. **e**, **f**, The numbers of subgraphs are obtained. **g**, **h**, The nodes with highest centrality of each subgraph are highlighted. **i**, **j**, After removing the nodes without highlighting, the density of subgraphs is calculated.

## References

1. Bar-On, Y. M., Phillips, R. & Milo, R. The biomass distribution on Earth. Proc. Natl. Acad. Sci. USA 115, 6506–6511 (2018).

2. Evert, R. F. *Esau’s Plant Anatomy: Meristems, Cells, and Tissues of the Plant Body: Their Structure, Function, and Development*. (John Wiley & Sons, 2006).

3. Cronk, Q. C. B. & Forest, F. Comparative and Evolutionary Genomics of Angiosperm Trees (eds Andrew Groover & Quentin Cronk) 1–17 (Springer International Publishing, 2017).

4. Strijk, J. S., Hinsinger, D. D., Zhang, F. & Cao, K. *Trochodendron aralioides*, the first chromosome-level draft genome in Trochodendrales and a valuable resource for basal eudicot research. GigaScience 8, giz136 (2019).

5. Magallón, S., Gómez-Acevedo, S., Sánchez-Reyes, L. L. & Hernández-Hernández, T. A metacalibrated time-tree documents the early rise of flowering plant phylogenetic diversity. New Phytol. 207, 437–453 (2015).

6. Greb, T. & Lohmann, J. U. Plant stem cells. Curr. Biol. 26, R816–821 (2016).

7. Bossinger, G. & Spokevicius, A. V. Sector analysis reveals patterns of cambium differentiation in poplar stems. J. Exp. Bot. 69, 4339–4348 (2018).

8. Johnsson, C. & Fischer, U. Cambial stem cells and their niche. Plant Sci. 252, 239– 245 (2016).

9. Shi, D., Lebovka, I., López-Salmerón, V., Sanchez, P. & Greb, T. Bifacial cambium stem cells generate xylem and phloem during radial plant growth. Development 146, dev171355 (2019).

10. Smetana, O. et al. High levels of auxin signalling define the stem-cell organizer of the vascular cambium. Nature 565, 485–489 (2019).

11. Denyer, T. et al. Spatiotemporal developmental trajectories in the *Arabidopsis* root revealed using high-throughput single-cell RNA sequencing. Dev. Cell 48, 840– 852 (2019).

12. Jean-Baptiste, K. et al. Dynamics of gene expression in single root cells of *Arabidopsis thaliana*. Plant Cell 31, 993–1011 (2019).

13. Ryu, K. H., Huang, L., Kang, H. M. & Schiefelbein, J. Single-cell RNA sequencing resolves molecular relationships among individual plant cells. Plant Physiol. 179, 1444–1456 (2019).

14. Shulse, C. N. et al. High-throughput single-cell transcriptome profiling of plant cell types. Cell Rep. 27, 2241–2247 (2019).

15. Turco, G. M. et al. Molecular mechanisms driving switch behavior in xylem cell differentiation. Cell Rep. 28, 342–351 (2019).

16. Wendrich, J. R. et al. Vascular transcription factors guide plant epidermal responses to limiting phosphate conditions. Science 370, 810 (2020).

17. Kim, J.-Y. et al. Distinct identities of leaf phloem cells revealed by single cell transcriptomics. Plant Cell 33, 511–530 (2021).

18. Xu, X. et al. Single-cell RNA sequencing of developing maize ears facilitates functional analysis and trait candidate gene discovery. Dev. Cell 56, 557–568 (2021).

19. Liu, Q. et al. Transcriptional landscape of rice roots at the single-cell resolution. Mol. Plant 14, 384–394 (2021).

20. Zhang, T. Q., Chen, Y. & Wang, J. W. A single-cell analysis of the *Arabidopsis* vegetative shoot apex. Dev. Cell 56, 1056–1074 (2021).

21. Zhang, T. Q., Chen, Y., Liu, Y., Lin, W. H. & Wang, J. W. Single-cell transcriptome atlas and chromatin accessibility landscape reveal differentiation trajectories in the rice root. Nat. Commun. 12, 2053 (2021).

22. Lopez-Anido, C. B. et al. Single-cell resolution of lineage trajectories in the Arabidopsis stomatal lineage and developing leaf. Dev. Cell 56, 1043–1055 (2021).

23. Larson, P. R. The Vascular Cambium*: Development and Structure*. (Springer Science & Business Media, 2012).

24. Smith, R. A. et al. Neighboring parenchyma cells contribute to *Arabidopsis* xylem lignification, while lignification of interfascicular fibers is cell autonomous. Plant Cell 25, 3988–3999 (2013).

25. Smith, R. A. et al. Defining the diverse cell populations contributing to lignification in *Arabidopsis* stems. Plant Physiol. 174, 1028–1036 (2017).

26. Blokhina, O. et al. Ray parenchymal cells contribute to lignification of tracheids in developing xylem of Norway spruce. Plant Physiol. 181, 1552–1572 (2019).

27. Pesquet, E. et al. Non-cell-autonomous postmortem lignification of tracheary elements in *Zinnia elegans*. Plant Cell 25, 1314–1328 (2013).

28. Barros, J., Serk, H., Granlund, I. & Pesquet, E. The cell biology of lignification in higher plants. Ann. Bot. 115, 1053–1074 (2015).

29. McQueen-Mason, S., Durachko, D. M. & Cosgrove, D. J. Two endogenous proteins that induce cell wall extension in plants. Plant Cell 4, 1425–1433 (1992).

30. Cho, H. T. & Cosgrove, D. J. Altered expression of expansin modulates leaf growth and pedicel abscission in *Arabidopsis thaliana*. Proc. Natl. Acad. Sci. USA 97, 9783–9788 (2000).

31. Scott, D. G. On the distribution of chlorophyll in the young shoots of woody plants. Ann. Bot. **os****-**21, 437–439 (1907).

32. Berveiller, D., Kierzkowski, D. & Damesin, C. Interspecific variability of stem photosynthesis among tree species. Tree Physiol. 27, 53–61 (2007).

33. Avila, E., Herrera, A. & Tezara, W. Contribution of stem CO_2_ fixation to whole-plant carbon balance in nonsucculent species. Photosynthetica 52, 3–15 (2014).

34. De Roo, L., Salomón, R. L., Oleksyn, J. & Steppe, K. Woody tissue photosynthesis delays drought stress in *Populus tremula* trees and maintains starch reserves in branch xylem tissues. New Phytol. 228, 70–81 (2020).

35. Chiang, M.-H. & Greb, T. How to organize bidirectional tissue production? Curr. Opin. in Plant Biol. 51, 15–21 (2019).

36. Zhu, Y., Song, D., Xu, P., Sun, J. & Li, L. A *HD-ZIP III* gene, *PtrHB4*, is required for interfascicular cambium development in *Populus*. Plant Biotechnol. J. 16, 808– 817 (2018).

37. Sundell, D. et al. AspWood: high-spatial-resolution transcriptome profiles reveal uncharacterized modularity of wood formation in *Populus tremula*. Plant cell 29, 1585–1604 (2017).

38. Barghoorn, E. S. The ontogenetic development and phylogenetic specialization of rays in the xylem of dicotyledons. I. the primitive ray structure. Am. J. Bot. 27, 918–928 (1940).

39. Braun, H. J. Beiträge zur Entwicklungsgeschichte der Markstrahlen. Botan. Stud. 4, 73–131 (1955).

40. Xu, B. et al. Contribution of NAC transcription factors to plant adaptation to land. Science 343, 1505 (2014).

41. Baran, Y. et al. MetaCell: analysis of single-cell RNA-seq data using K-nn graph partitions. Genome Biol. 20, 206 (2019).

42. Ohtani, M. et al. A NAC domain protein family contributing to the regulation of wood formation in poplar. Plant J. 67, 499–512 (2011).

43. Liu, B. et al. Transcriptional reprogramming of xylem cell wall biosynthesis in tension wood. Plant Physiol. 186, 250–269 (2021).

44. Reid, A. J. et al. Single-cell RNA-seq reveals hidden transcriptional variation in malaria parasites. eLife 7, e33105 (2018).

45. Liang, C. B., Baas, P., Wheeler, E. A. & Shuming, W. Wood anatomy of trees and shrubs from China. VI. magnoliaceae. IAWA Journal 14, 391–412 (1993).

46. Hao, Y. et al. Integrated analysis of multimodal single-cell data. Cell 184, 3573– 3587 (2021).

47. Bollhöner, B., Prestele, J. & Tuominen, H. Xylem cell death: emerging understanding of regulation and function. J. Exp. Bot. 63, 1081–1094 (2012).

48. Strabala, T. J. & MacMillan, C. P. The *Arabidopsis* wood model—the case for the inflorescence stem. Plant Sci. 210, 193–205 (2013).

49. Chaffey, N., Cholewa, E., Regan, S. & Sundberg, B. Secondary xylem development in *Arabidopsis*: a model for wood formation. Physiol. Plant. 114, 594–600 (2002).

50. Nieminen, K. M., Kauppinen, L. & Helariutta, Y. A weed for wood? *Arabidopsis* as a genetic model for xylem development. Plant Physiol. 135, 653–659 (2004).

51. Lin, Y. C. et al. SND1 transcription factor-directed quantitative functional hierarchical genetic regulatory network in wood formation in *Populus trichocarpa*. Plant Cell 25, 4324–4341 (2013).

52. Lin, Y. C. et al. A simple improved-throughput xylem protoplast system for studying wood formation. Nat. Protoc. 9, 2194–2205 (2014).

53. Yeh, C. S. et al. A novel synthetic-genetic-array–based yeast one-hybrid system for high discovery rate and short processing time. Genome Res. 29, 1343–1351 (2019).

54. Keren-Shaul, H. et al. MARS-seq2.0: an experimental and analytical pipeline for indexed sorting combined with single-cell RNA sequencing. Nat. Protoc. 14, 1841–1862 (2019).

55. Yu, D., Huber, W. & Vitek, O. Shrinkage estimation of dispersion in negative binomial models for RNA-seq experiments with small sample size. Bioinformatics 29, 1275–1282 (2013).

56. Robinson, M. D. & Smyth, G. K. Moderated statistical tests for assessing differences in tag abundance. Bioinformatics 23, 2881–2887 (2007).

57. Benjamini, Y. & Hochberg, Y. Controlling the false discovery rate: a practical and powerful approach to multiple testing. J. R. Stat. Soc. Series B Stat. Methodol. 57, 289–300 (1995).

58. Chen, S., Zhou, Y., Chen, Y. & Gu, J. fastp: an ultra-fast all-in-one FASTQ preprocessor. Bioinformatics 34, i884–i890 (2018).

59. Kim, D., Langmead, B. & Salzberg, S. L. HISAT: a fast spliced aligner with low memory requirements. Nat. Methods 12, 357–360 (2015).

60. Kim, D., Paggi, J. M., Park, C., Bennett, C. & Salzberg, S. L. Graph-based genome alignment and genotyping with HISAT2 and HISAT-genotype. Nat. Biotechnol. 37, 907–915 (2019).

61. Pertea, M. et al. StringTie enables improved reconstruction of a transcriptome from RNA-seq reads. Nat. Biotechnol. 33, 290–295 (2015).

62. Love, M. I., Huber, W. & Anders, S. Moderated estimation of fold change and dispersion for RNA-seq data with DESeq2. Genome Biol. 15, 550 (2014).

63. Shi, R. et al. Tissue and cell-type co-expression networks of transcription factors and wood component genes in *Populus trichocarpa*. Planta 245, 927–938 (2017).

64. Altschul, S. F. et al. Gapped BLAST and PSI-BLAST: a new generation of protein database search programs. Nucleic Acids Res. 25, 3389–3402 (1997).

65. Camacho, C., et al. BLAST+: architecture and applications. BMC Bioinformatics 10, 421 (2009).

66. Li, L., Stoeckert, C. J., Jr. & Roos, D. S. OrthoMCL: identification of ortholog groups for eukaryotic genomes. Genome Res. 13, 2178–2189 (2003).

67. Enright, A. J., Van Dongen, S. & Ouzounis, C. A. An efficient algorithm for large-scale detection of protein families. Nucleic Acids Res. 30, 1575–1584 (2002).

68. Stuart, T. et al. Comprehensive integration of single-cell data. Cell 177, 1888–1902 (2019).

69. Friedman, J. H. & Rafsky, L. C. Multivariate generalizations of the Wald-Wolfowitz and Smirnov two-sample tests. Ann. Stat. 7, 697–717 (1979).

70. Chen, H., Chen, X. & Su, Y. A weighted edge-count two-sample test for multivariate and object data. J. Am. Stat. Assoc. 113, 1146–1155 (2018).

71. Csardi, G. & Nepusz, T. The igraph software package for complex network research. Int. J. Complex Syst. 1695, 1–9 (2006).

72. Venables, W. N. & Ripley, B. D. *Modern Applied Statistics with S* (eds W. N. Venables & B. D. Ripley) 107–138 (Springer New York, 2002).

73. Street, K. et al. Slingshot: cell lineage and pseudotime inference for single-cell transcriptomics. BMC Genomics 19, 477 (2018).

74. Finak, G. et al. MAST: a flexible statistical framework for assessing transcriptional changes and characterizing heterogeneity in single-cell RNA sequencing data. Genome Biol. 16, 278 (2015).

